# The mechanical response of vinculin

**DOI:** 10.1101/2023.05.25.542235

**Authors:** Xuyao Liu, Yinan Wang, Mingxi Yao, Karen B. Baker, Benjamin Klapholz, Nicholas H. Brown, Benjamin T. Goult, Jie Yan

## Abstract

Vinculin is a mechanosensitive adapter protein that links the actin network to cell-extracellular matrix adhesions and cell-cell adhesions. It is perhaps the best characterized mechanoeffector, as it is recruited to sites of adhesion in response to force on the mechanotransducers talin and alpha-catenin. Here we examined the mechanical properties of vinculin to assess its potential role as a mechanotransducer. We find that at physiological loading rates, the structural domains of vinculin unfold at forces in the 5-15 pN range and rapidly refold when forces are reduced back to 1 pN. Thus, vinculin domains also have the potential to act as force dependent molecular switches, akin to those in talin and alpha-catenin. As with the force dependent switches in talin, the unfolding of these domains in vinculin introduces large extension changes in the vinculin cytoskeletal linkage up to 150 nm with 20-30 nm steps of unfolding. Modelling of the tension-dependent interactions of the unstructured vinculin linker region with a model protein containing two SH3 domains indicated that even unstructured protein regions can mediate force-dependent interactions with ligands, where the binding of a dual-SH3 model protein is predicted to be significantly suppressed by forces greater than 10 pN. Together, these findings suggest that vinculin has a complex mechanical response with force-dependent interaction sites, suggesting it also acts as a mechanotransducer, recruiting partners in response to force.

## I. INTRODUCTION

In recent years, there has been growing appreciation that cells not only respond to the chemical signals in their environment, but also to mechanical signals, such as the stiffness of the tissue and surrounding extracellular matrix [1, 2]. Sensing the mechanical environment requires mechanotransducers: molecules that can convert mechanical forces into chemical changes within the cell. A current paradigm for mechanotransduction is provided by the mechanotransducers talin and alpha-catenin, which recruit the mechanoeffector vinculin in a force-dependent manner [3-5].

The force-dependent recruitment of vinculin requires the mechanical unfolding of domains within talin and alpha-catenin, key cytoskeletal linkers at the adhesive junctions mediating cell-extracellular matrix and cell-cell adhesion respectively [6, 7]. This occurs because the vinculin binding sites (VBSs) are buried within the folded alpha-helical bundles, and are exposed by force unfolding these domains. Thus, these domains function as mechanical switches, as they bind some proteins in their folded state, and others in their unfolded state [3]. Vinculin is one of the best characterized mechanoeffectors, as its recruitment to sites of cell adhesion is force dependent.

The interaction of vinculin with talin and alpha-catenin is not simply regulated by the exposure of VBSs as vinculin itself can also adopt closed and open conformations, altering its interactions with its binding partners. Vinculin consists of a number of bundles of four alpha-helices. The number of these bundles varies from 6-8, depending on the species, and in some cases pairs of bundles share an extended alpha-helix, thereby forming a domain. Thus, vertebrate vinculin consists of domains D1 (bundles 1 and 2), D2 (bundles 3 and 4), D3 (bundles 5 and 6), D4 (bundle 7), an unstructured proline-rich linker and a conserved 5-helix bundle, vinculin tail, Vt (bundle 8) at the C-terminus. The major interactions with vinculin reported to date involve D1, which binds VBS, the linker, which binds SH3 domain-containing partners such as CAP and p130Cas [8], and Vt which binds actin. Alpha-catenin is a paralog of vinculin, with a similar structure, and the bulk of talin is also formed of helical bundles, some four-helix bundles and the majority with the addition of a fifth helix [9]. Inter-domain interactions within the vinculin head organize it into a pincer-like structure [10, 11] (Fig. 1a).

**Fig. 1.**
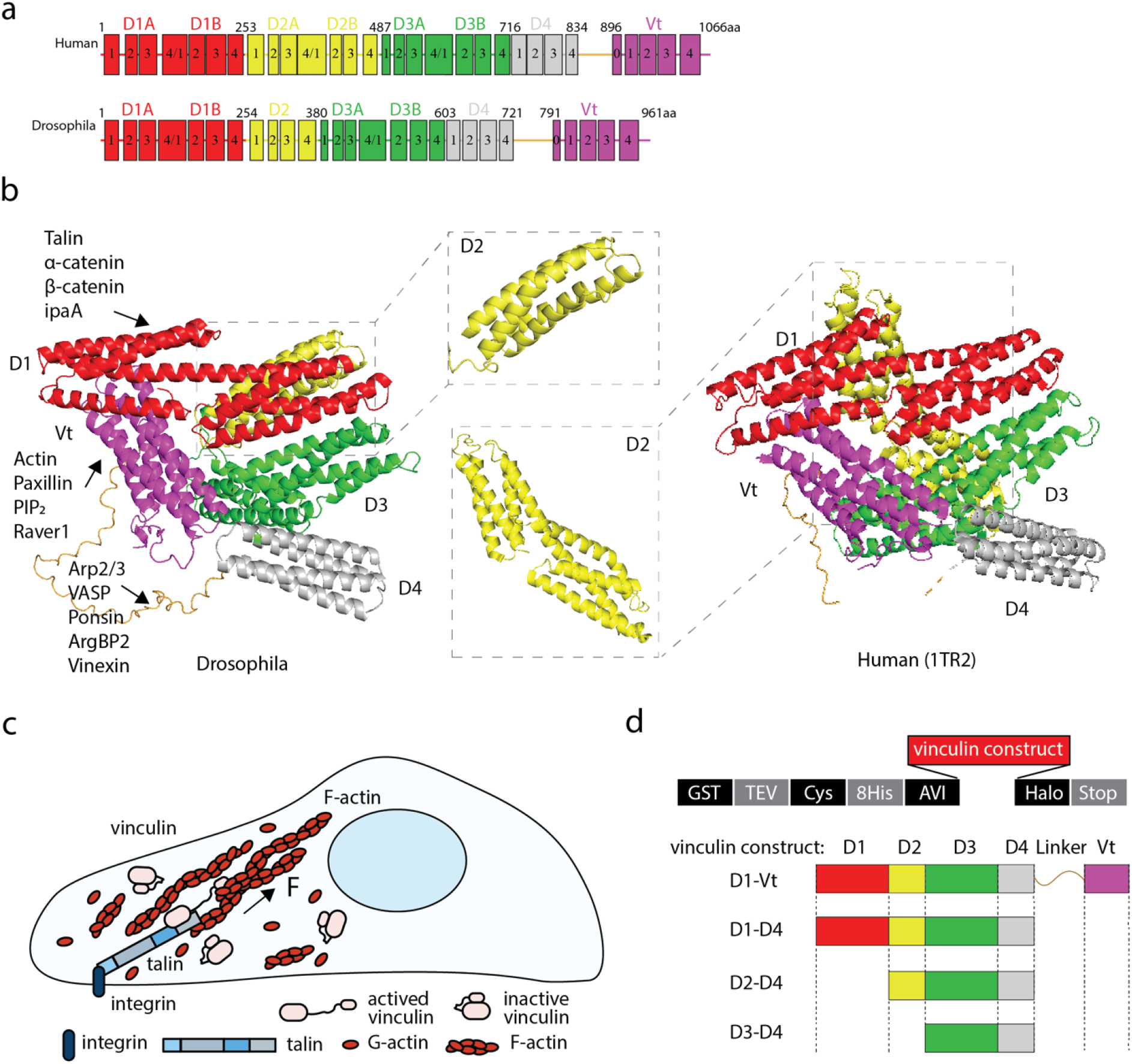
Stretching vinculin. (a) Domain structures of Drosophila and human vinculin. (b) Structural model of Drosophila vinculin (dVinc) predicted by AlphaFold2 in an auto-inhibited configuration (D1 in red, D2 in yellow, D3 in green, D4 in grey and Vt in purple with linker in gold). Right. The structure of full-length human vinculin (hVinc) (PDB ID. 1TR2). Inset. The D2 domain is shown enlarged with a single 4-helix bundle in dVinc and a double 4-helix bundle in hVinc. The known vinculin binding proteins are listed for individual vinculin domains. (c) Illustration of the force-transmission supramolecular linkage between integrins engaging the extracellular matrix and the actin network mediated by talin (grey/blue) and vinculin (pink). The talin-bound, activated vinculin mediates force transmission from talin to actin. (d) The vector design used in these experiments is shown where a GST-tag is used for purification and once cleaved using TEV protease yields a protein of interest with N-terminal AVI-tag and C-terminal Halo-tag for tethering to allow stretching. Four vinculin constructs were designed: D1-Vt (full-length vinculin), D1-D4, D2-D4 and D3-D4.

Vinculin exists in equilibrium between closed and open conformations. In the closed conformation it does not bind VBSs or actin [12] due to an autoinhibitory interaction between the head and tail domains [13, 14]. The open conformation is able to engage with talin/alpha-catenin and actin, and is stabilized by these interactions [15, 16]. Force strengthens these interactions, as observed in cells [17]. When force is reduced, the mechanosensitive interactions weaken and vinculin disengages and returns to its closed conformation. The importance of this regulatory mechanism was highlighted by the observation that in Drosophila, expression of constitutively open vinculin is more detrimental than removing vinculin [18].

Vinculin binding to actin enables its role in FA-mediated mechanotransduction [12, 19-24] and downregulation of vinculin leads to disruptions in cell rigidity sensing, cell adhesion, cell growth and migration, which in turn contribute to cancer metastasis [25-27]. Loss of vinculin has also been found to break the balance in skin stem cell differentiation [28], and has been linked to both dilated and hypertrophic cardiomyopathies in patients [29-31].

Active vinculin thus forms a mechanical linkage between the VBS-binding D1 domain at one end and the actin-binding Vt domain at the other end (Fig. 1b). As a result, actomyosin contraction will exert force on the other domains of vinculin. The key question we wished to address in this paper is whether in addition to being a key mechanoeffector of the mechanotransducing proteins talin and alpha-catenin, could vinculin also be a mechanotransducer, regulating the binding to its partners in a force dependent manner? To gain insights into this question, it is essential to quantify the mechanical responses of the tension-bearing domains and linker regions of vinculin. Based on these responses, we can predict how tension might influence the binding of factors associated with vinculin.

In this work, we investigated the mechanical response of full-length vinculin and the binding of a dual-SH3 model protein to the vinculin linker using an integrated approach combining single-molecule manipulation experiments, AlphaFold2 structural modelling and binding predictions, and modelling based on classic polymer theory and statistical physics. Our results reveal the complex mechanical response of vinculin and identify that the helical bundles in vinculin exhibit switch-like behavior. We experimentally show that 1) the force-bearing vinculin domains undergo dynamic unfolding and refolding structural transitions over physiological range of stretching conditions, 2) the dimensions of the vinculin molecule change dramatically as these domains unfold and refold. Further, results from structural and theoretical modelling suggest that 3) the vinculin linker is less accessible to binding factors in the auto-inhibited conformation, 4) tensile force of a few pN can extended vinculin and significantly change the binding affinity between the vinculin linker and its binding factors. Together, this work identifies vinculin as a complex mechanosensor and mechanotransducer.

## II. RESULTS

### Mechanical stability of vinculin domains

Our previous work showed that the different domains of the talin rod unfold at different forces, and rapidly refold when force is released [32]. We wished to explore whether vinculin shares these properties, and so could also function as a mechanotransducer. Our recent work benefited from the genetic tractability of Drosophila for the study of vinculin [18], so for this study we chose to initiate our single molecule studies with Drosophila vinculin. Drosophila vinculin has 7 helical bundles (Fig. 1a) and functions at room temperature, as the optimum temperature for Drosophila culture is 22-25°C. As the structure of Drosophila vinculin has not been solved, we predicted it using AlphaFold2 [33, 34], which produced a structure consistent with the structure previously modelled using MODBASE (ref. no. O46037, Supp. Fig. 1) based on the chicken vinculin crystal structure (PDB ID. 1ST6 [10]). The predicted structure shows high similarity with that of chicken (PDB IDs. 1ST6 [10] and 6NR7 [35]) and human (PDB ID. 1TR2 [9]) vinculin (Fig. 1a,b) and also with the vinculin homolog from sponge (PDB ID. 6BFI [36]). The main difference between the vinculins from different species is found in the D2 domain (Fig. 1a and 1b insets): the D2 domain in human and chicken vinculin contains two four-helix bundles connected by a long, centrally shared *α*-helix, the D2 domain in Drosophila vinculin is a single four-helix bundle, and D2 is absent entirely from sponge vinculin [36]. The tail domain (Vt) interacts with the head domains, resulting in a compact, lowest-energy autoinhibited, closed conformation. When activated, the open form of vinculin links talin to F-actin in the integrin-mediated mechanotransduction-pathway and force is exerted on vinculin (Fig. 1c). Vinculin also interacts with various cellular proteins as indicated (Fig. 1b) [6, 21, 37-41].

To understand the mechanical stability of the tension-bearing vinculin domains, we expressed segments of vinculin with suitable tags for stretching (Fig. 1d).

### The mechanical response of full-length vinculin

The mechanical stability of Drosophila full-length vinculin was investigated at room temperature (24°C) using an in-house constructed magnetic-tweezer setup [42, 43] (Fig. 2a). The height of the end-attached 2.8 *μ*m-diameter superparamagnetic bead (Dynabeads™ M-270, Invitrogen) was recorded as the force was increased at selected force loading rate. After each force-loading scan, the force was reduced to 1 pN for 30 s, allowing the unfolded domains to refold, and then the force-loading scan was repeated. Figure 2b shows representative force-height curves during repeating force-loading of a tether containing a full-length vinculin at a loading rate of 2.0±0.2 pN/s. The stepwise bead height increases indicate domain unfolding events. Most of the domains unfold within 7-15 pN. The data reveal a complex mechanical response, with 6-7 unfolding steps spread over the force range. The number of unfolding steps is consistent with the number of helical bundles (D1A, D1B, D2, D3A, D3B, D4, and Vt). The size of ∼ 30 nm of each extension step is also consistent with unfolding of one helical bundle. The data did not reveal a mechanically stable head-tail association, which would have manifested as an additional stepwise bead height increase (further discussed in the Discussion section).

**Fig. 2.**
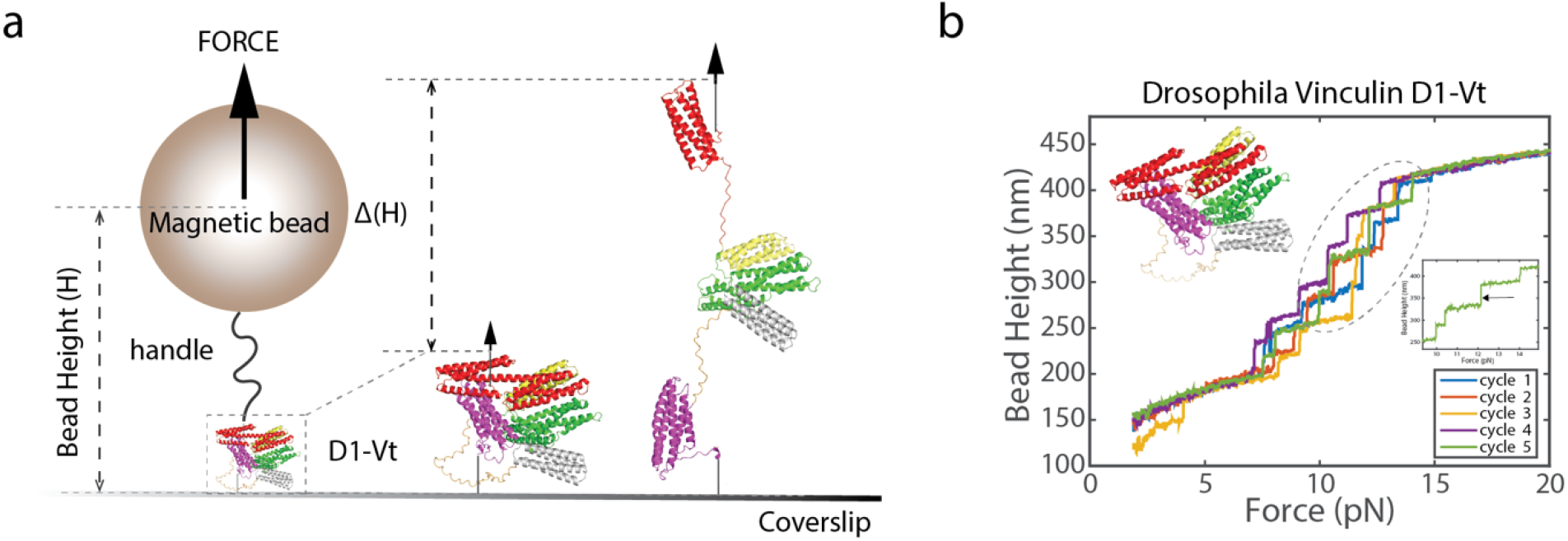
Mechanical response of full-length vinculin. (a) Full-length vinculin (D1-Vt shown in Fig. 1b,d) is tethered between coverslip and a superparamagnetic bead using a 572-bp DNA handle. Force is exerted on the bead and onto the protein resulting in a change in bead height (ΔH), domain unfolding events result in stepwise jumps in bead height. (b) Five representative consecutive force-height curves obtained from a tether at a loading rate of 2.0±0.2 pN/s. (Inset top left) structural model of full-length Drosophila vinculin. (Inset right) All of the bundles in vinculin are 4-helical except the Vinculin tail domain, which is a 5-helix bundle making its unfolding event readily identifiable by the larger step that results when it unfolds, indicated by the arrow.

We next examined the mechanical responses of tension-bearing subdomains in Drosophila vinculin. In these studies, we reduced the loading rate to 0.40±0.04 pN/s to extend the duration, allowing us to capture more detailed information on the dynamic unfolding and folding fluctuations of these subdomains.

### D3-D4 domains

The D4 domain is a single four-helix bundle and the D3 domain contains two four-helix bundles connected by a shared *α*-helix. Three distinct unfolding steps over a force range of 7-15 pN were observed in each scan (Fig. 3a, left) occurring over a range of forces. This indicates that the three *α*-helix bundles unfold independently. In each cycle, the unfolded D3 and D4 domains refolded with near 100% probability when the force was reduced to 1 pN and held for 30 s, enabling us to perform many mechanical unfolding and refolding cycles of the same domains before breakage.

**Fig. 3.**
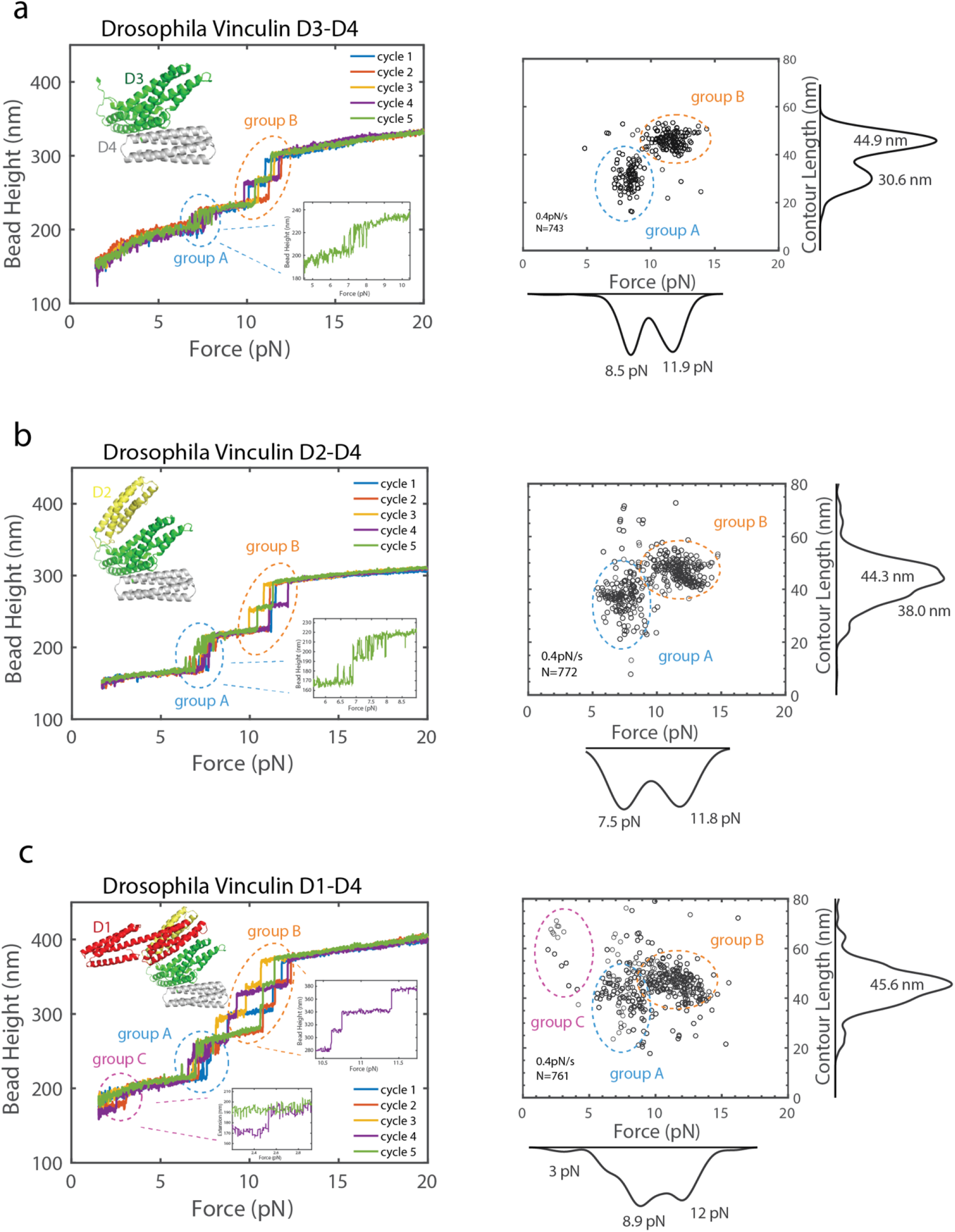
Mechanical responses of vinculin D3-D4, D2-D4 and D1-D4 sub-segments. (a-c) Five representative, consecutive force-height curves of the unfolding of vinculin D3-D4 (a), D2-D4 (b) and D1-D4 domains (c) at 0.40±0.04 pN/s loading rate are shown. The structural models of each sub-segment are shown in the upper left. For each sub-segment, the unfolding events are divided into groups based on the contour lengths of released polypeptide polymer and unfolding forces (right panels). The rapid fluctuations for group A unfolding events are shown enlarged as inset in D3-D4 (a) and D2-D4 domains (b). Two equilibrium unfolding/refolding fluctuation in group A are noted in D2-D4 (b). In the D1-D4 force-extension curves (c), two regions are enlarged corresponding to two unfolding signals from the two 4-helix bundles of the D1 domain. For each sub-segment, data points are obtained from >700 unfolding events on at least 4 independent tethers.

We next converted the force-dependent step sizes into the contour length of protein peptide released after unfolding (Supp. Info. 1 & Supp. Fig. 2) to generate a force-contour length scatter plot (Fig. 3a, right). This plot includes more than 700 unfolding events from 13 independent tethers and shows two clear groups (group A and group B) with differential unfolding forces (Supp. Fig. 3). Group A contains the weakest *α* -helical bundle which undergoes rapid reversible unfolding and refolding transitions (Fig. 3a, left) at forces around 8 pN. Similar rapid unfolding/refolding fluctuations were seen previously for the weakest domain in talin, R3 [32] when at ∼5 pN of force. In contrast, the other two *α*-helical bundles (group B) unfold at forces of 10-15 pN and do not refold at this force, only refolding once force is released at the end of the cycle. This mechanical hysteresis property of the domains was also similar to what was seen in the other talin rod domains [32]. As the contour lengths are dependent on the primary sequence of the domain, we could calculate that the group B data in Fig. 3a corresponds to the unfolding of D4 and the larger sub-domain from D3, D3B. The group A corresponds to the unfolding of the smaller sub-domain in D3, D3A (Supp. Table 1 & Supp. Fig. 4).

**Fig. 4.**
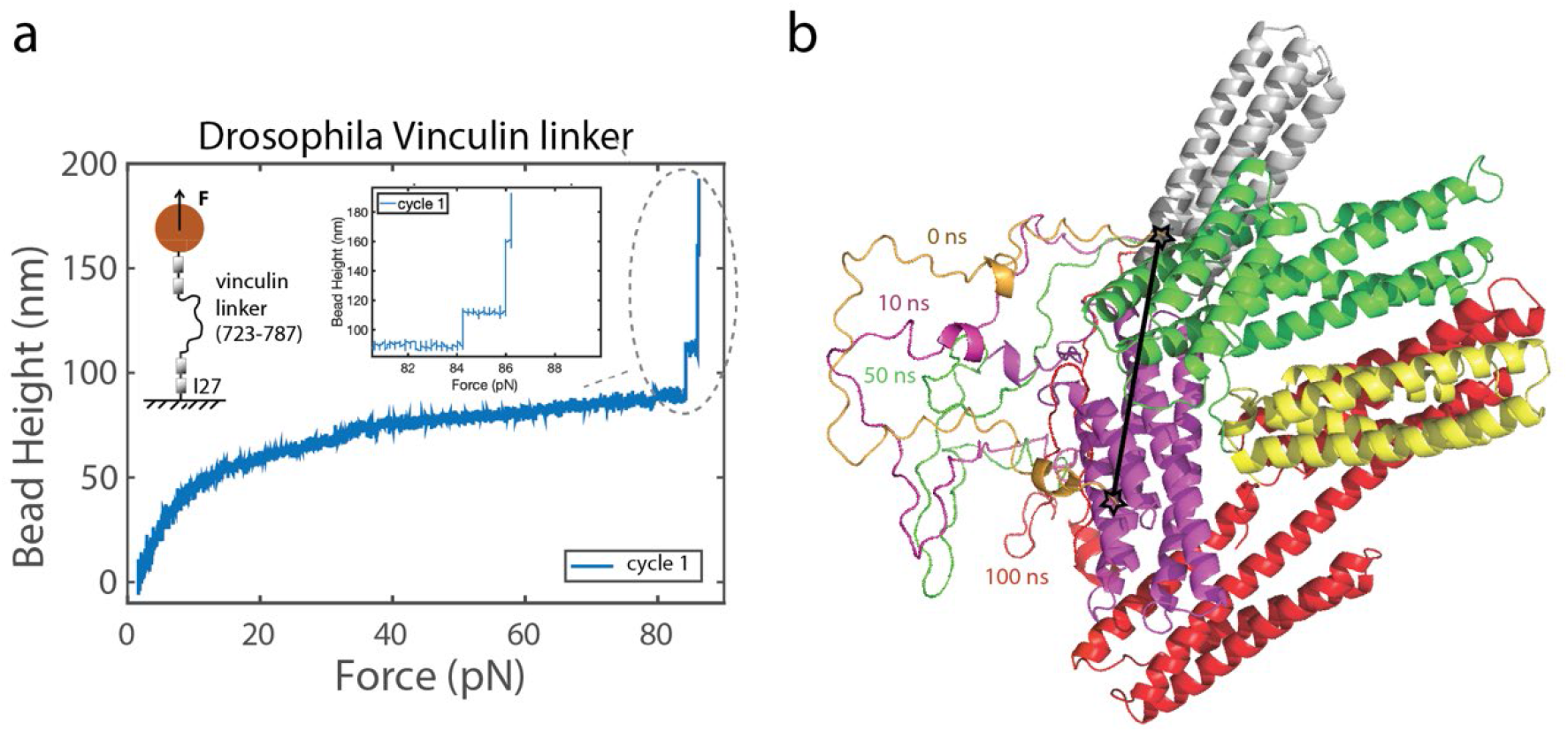
Vinculin linker is an unstructured polypeptide. (a) A representative consecutive force-height curve of a tether containing the vinculin linker is shown. The linker is spanned between two titin I27 domains on each side (inset). The signature unfolding steps of I27 domains serves as a specific control of tether. No unfolding signals were observed from the linker. (b) Snapshots of Drosophila vinculin linker conformations when relaxed in molecular dynamics (MD) simulation. The initial conformation of the linker (gold) was extracted from the full-length vinculin predicted by AlphaFold2, followed by MD simulation with a fixed end-to-end distance of 4.63 nm for up to 100 ns. The snapshots of the linker conformations extracted at different time points were then added back to the vinculin structure with the same position of the ends (denoted by stars at the ends of the black line) on D4 and Vt.

### D2-D4 domains

Having quantified the mechanical response of the D3-D4 domains, we next extended the experiment to include the additional four-helix bundle in the D2 domain. The left panel of Fig. 3b shows the representative force-bead height curves obtained during force-increase scans at the same loading rate of 0.4 pN/s and the force-contour length scatter plot is shown accordingly (Fig. 3b, right). When compared to the data from D3-D4, the signature of the additional D2 domain is clear. D2 also undergoes rapid reversible unfolding and refolding transitions at forces around 7-8 pN. Therefore, the D2 domain is assigned to the mechanical group A.

### D1-D4 domains

The results from similar experiments performed for the construct D1-D4 are shown in Fig. 3c. Compared to data obtained from D2-D4, the mechanical signature from D1 is revealed. Two distinct signals from D1 are evident, one contributing to an additional unfolding signal in the mechanical group B, however, D1 domain also contributes to a very mechanically unstable unfolding event, with an unfolding signal at forces around 2-3 pN at the loading rate (group C, Fig. 3c). The group C unfolding signal was not observed in every force-increase scan, indicating that the corresponding domain did not always successfully refold when the tether was held at the refolding force (1.0 ± 0.1 pN) over 30 seconds (Supp. Fig. 5). The appearance of two distinct unfolding signatures of D1 is consistent with the existence of two *α*-helical bundles, D1A and D1B, in D1. D1A is responsible for the binding to vinculin binding sites, and D1B is a force-bearing *α* -helical bundle in vinculin-mediated force-transmission pathway. Due to the similar sizes of D1A and D1B, we cannot assign them into the respective groups.

**Fig. 5.**
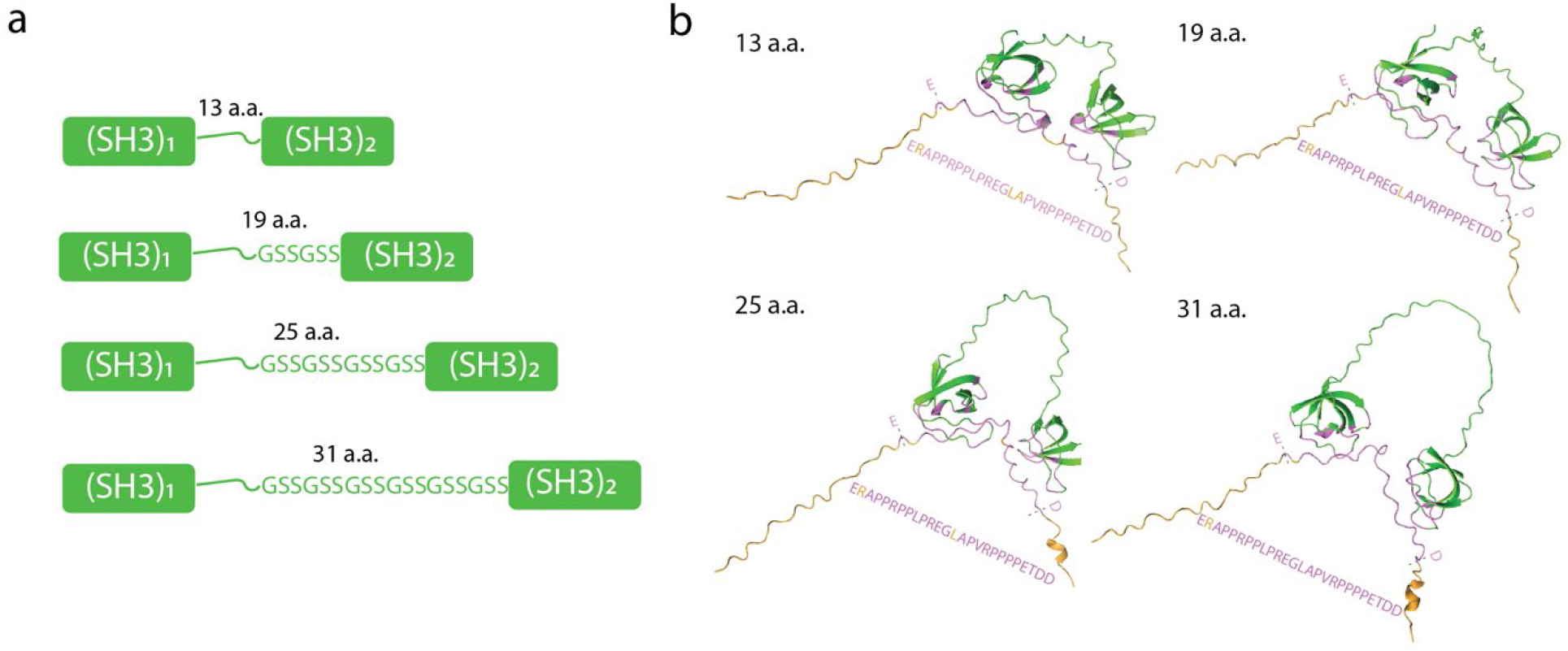
Predicted structural interactions between the vinculin linker and SH3 repeats separated by various inter-SH3 lengths. (a) Illustrations of SH3 repeats with 13 a.a., 19 a.a., 25 a.a., and 31 a.a. inter-SH3 length by adding repeats of GSSGSS. (b) Predicted structural complexes of the isolated vinculin linker (yellow) and SH3 repeats with various inter-SH3 lengths (green). The predicted interactions are highlighted in purple with the amino acid sequences shown.

Comparing D1-D4 (Fig. 3c) with full-length vinculin (Fig. 2b) we can identify the additional unfolding step which belongs to the Vt domain, which has a larger force-dependent step size expected from a five-helix bundle. Moreover, the observation that one of the subdomains, either D1A or D1B, did not refold in the majority of the force-loading cycles accounts for the presence of only six unfolding steps in four of the five cycles of stretching full-length vinculin (Fig. 2b).

Overall, the results in this section show that the force-bearing *α*-helical bundles in the vinculin head domains (D1-D4) show switch-like mechanical properties, unfolding independently over a force range from 2-15 pN at the loading rate of 0.4 pN/s. Among the force-bearing domains, we identify two distinct groups of mechanical behaviors, D4, D3B, and a sub-domain from D1 are more mechanically stable (group B). Each switch in group B exhibits mechanical hysteresis, and does not refold at the same force as it unfolds. The second group (group A) includes the mechanically weaker domains D2 and D3A which undergo rapid fluctuations in unfolding/refolding around 8 pN of force. One sub-domain in D1 does not fit well into either of these distinct groupings so we included a third group (group C). Notably, the extension of the mechanical linkage mediated by vinculin can change from around 10 nanometers (low force, relaxed conformation) to more than 150 nanometers at forces up to 15 pN when all force-bearing domains (D1B, D2, D3, and D4) are unfolded (Supp. Info. 2).

### The vinculin linker does not contain force-unfolding steps but is mechanosensitive

Besides the force-bearing vinculin domains, the 59 a.a. linker domain between the vinculin head and the tail also bears force. The linker region is not fully visible in the crystal structures solved to date and is predominantly unstructured. As the linkers mechanical response was not visible in our stretching of full-length vinculin (Fig. 2b) we designed an additional construct to allow us to visualize how the linker region responds to force. In this construct the vinculin linker was introduced between two pairs of titin I27 domains which provide a signature mechanical signal. Single-molecule stretching of the linker region did not show any unfolding steps during force-increase scans, except the signature unfolding steps from the I27 domains, which is consistent with the linker being an unstructured polypeptide polymer (Fig. 4a). This is also consistent with the prediction by AlphaFold2 that the vinculin linker is unstructured (Fig. 4b and Supp. Fig. 6).

**Fig. 6.**
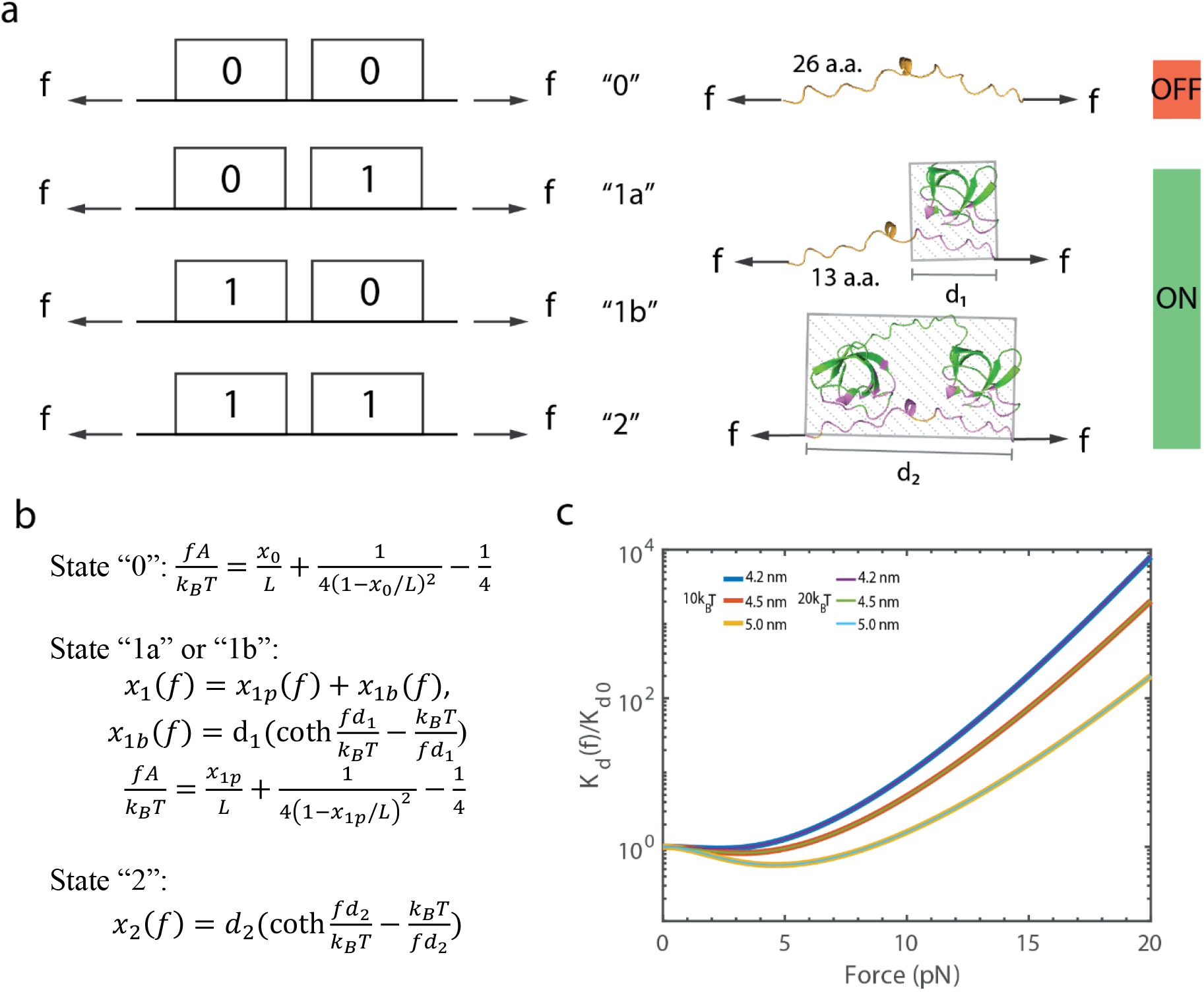
The model of the force-dependence of the vinculin linker engaging tandem SH3 domains. (a) A 26 amino acid (a.a.) segment of the vinculin linker is modeled in complex with SH3 repeats, as depicted in the schematic illustrations. The tandem interaction has four states: “0” for no binding, “1a” or “1b” for single binding, and “2” for dual binding. The OFF state indicates no binding, while the ON state denotes that one of the binding sites is bound. In state “0”, the linker is a flexible polymer; in state “1a” or “1b,” it is a combination of a flexible polymer region and a rigid bound region; and in state “2,” it adopts a rigid bound conformation. The linear distances between the force-attaching points on the rigid bound regions for states “1a/b” and “2” are denoted by *d*_1_ and *d*_2_, respectively. (b) The force-extension curves of the 26 amino acid linker region for each state are as follows. In state “0,” the force-extension curve is described by Marko-Siggia formula suitable for flexible polypeptide polymer with a contour length *L* and a bending persistence length *A* = 0.8 nm. In state “1a” or “1b,” the force-extension curve is a summation of the contributions from the unbound polymer region *x*_1*p*_(*f*) and the rigid bound region *x*_1*b*_(*f*) of a length *d*_1_. In state “2,” the force-extension curve is based on approximating the entire region as a rigid body of a length *d*_2_ (details in Supp. Info 3). (c) Force-dependent dissociation constant K_*d*_(*f*) is plotted against K_*d*0_ = K_*d*_(*f* = 0) for the single-SH3 binding energy of 10*k*_*B*_ *T* and 20*k*_*B*_ *T* with *d*_2_ values ranging from 4.2 to 5.0 nm.

The conformation of the vinculin linker in the autoinhibited, compact form of Drosophila vinculin is likely limited due to the entropic elasticity of an unstructured polypeptide polymer. AlphaFold2 predicts that the linker on the compact conformation of full-length vinculin has a fixed extension of 4.60±0.04 nm (mean±S.D.) (Supp. Table 2 & Supp. Fig. 7). Considering this constraint, the tensile force within the linker can be estimated to be around 3 pN (Supp. Fig. 8). This tension may pull the linker towards the structural domains (Fig. 4b, Supp. Movie 1), potentially leading to a less accessible conformation for binding factors, as binding would cause steric clashes with the nearby structured domains. This observation aligns with the full-atom molecular dynamics (MD) simulation of the linker over a 100 ns timeframe, beginning with the initial conformation taken from the complete vinculin structure as predicted by AlphaFold2 (Supp. Movie 1).

**Fig. 7.**
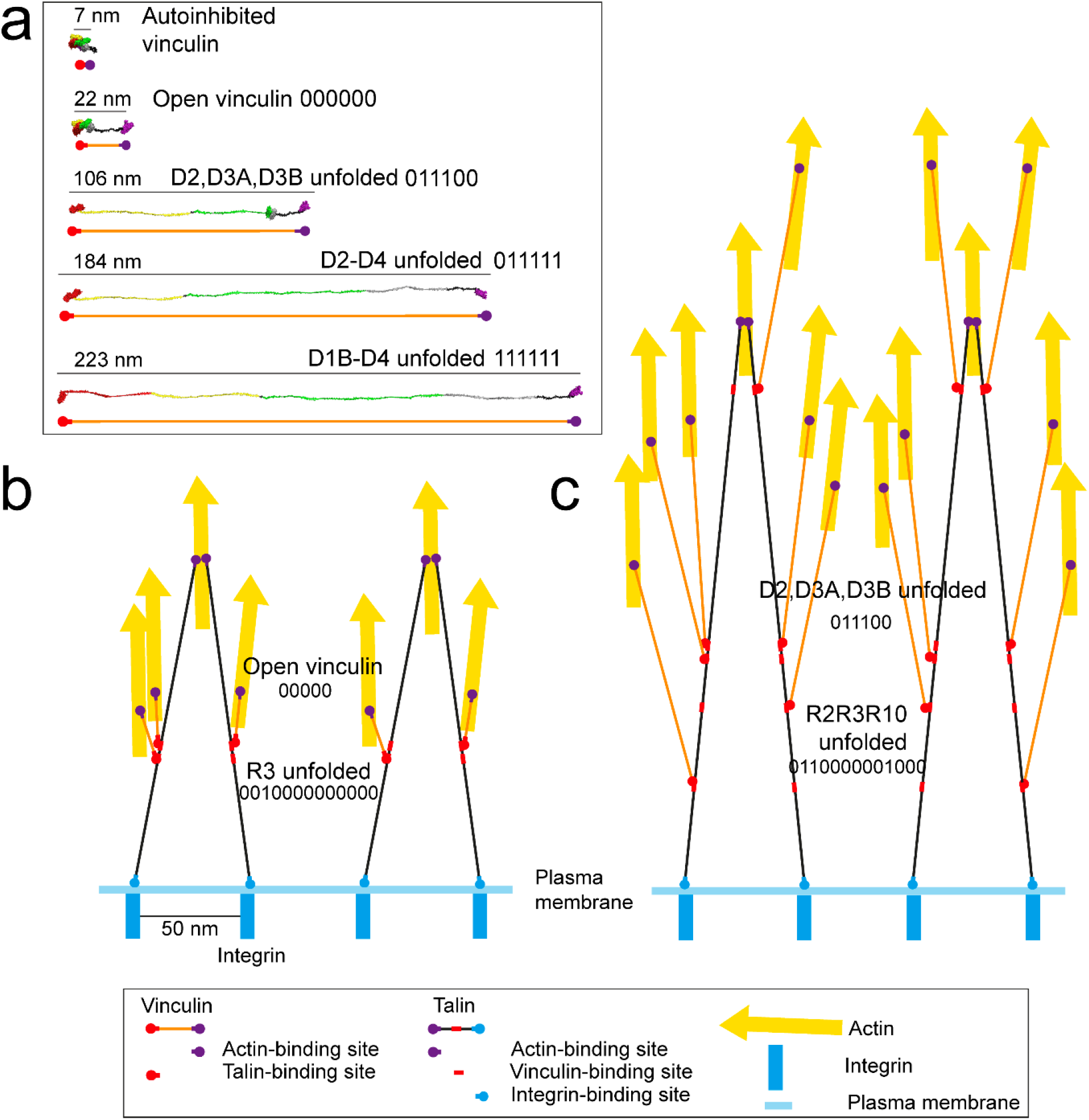
Vinculin to scale. (a) Structural model of vinculin in different states; autoinhibited, open, extended and fully extended to show the large differences in dimensions that occur. The six switch domains; D1B, D2A, D2B, D3A, D3B and D4 that are in the linkage between the talin-binding site in D1A and the actin-binding site in Vt are shown in different states. Having two states, folded and unfolded these domains can be represented as binary switches with states 0 (folded) and 1 unfolded). The state dependent contour lengths of vinculin are provided. (b-c) “To-scale” illustrations of a talin-vinculin-actin linkage in two different mechanical states. (b) With just one talin domain, R3, unfolded and with vinculin open but all domains folded (c) With three talin domains, R2, R3 and R10 unfolded, and the three weakest domains in vinculin, D2, D3A and D3B unfolded. The dimensions of the linkage and the distance between attachment sites can extend to 100’s of nanometers as a function of the switch patterns.

### Prediction of the complex formed between dual-SH3 repeats and vinculin linker based on AlphaFold2

The vinculin linker is a hotspot for ligand binding and contains four proline-rich motifs that engage with Src-homology 3 (SH3) domain containing proteins. Having shown that the disordered vinculin linker is forced in close proximity to the folded vinculin domains in the compact arrangement that presumably could suppress binding of ligands to the linker, we next wanted to theoretically explore whether force could modulate binding to the linker when the linker is fully exposed in the open conformation of vinculin. If the end-to-end extension of the linker is altered by the binding factor, then this will introduce a force dependence to the affinity of the interaction [44]. As many factors bind to the vinculin linker multivalently via their multiple SH3 domains [8, 39, 45, 46], we based our investigation on a dual-SH3 model protein consisting of the first two SH3 domains of Drosophila CAP (UniProtKB - Q966U7_DROME).

The conformation of the linker, when bound to its binding factor, is crucial for the theoretical modeling of force-dependent binding affinity. To predict the conformation of the vinculin linker (residues 722-786) bound to the dual-SH3 repeat, we utilized AlphaFold2. The structural modelling showed good agreement with the solution structures of the two SH3 domains bound to vinculin solved previously validating the approach [47]. Specifically, the 3^rd^ and 4^th^ proline-rich motifs which include the shared core 24 amino acid sequence, ERAPPRPPLPREGLAPVRPPPPET (residues 751-774) were encompassed within the bound complex with the dual-SH3 (Fig. 5b and Supp. Table 3).

Since the inter-SH3 length is a variable feature among different SH3-containing linker binding factors, we next performed predictions for dual-SH3 repeats with varying inter-SH3 lengths of 13, 19, 25 and 31 amino acids (Fig. 5a). In all cases, AlphaFold2 predicted the same interaction with the vinculin linker with high confidence (Fig. 5b, Supp. Table 4).

Although the AlphaFold2-based predictions suggest that the SH3 domain binding sites on the vinculin linker for the dual-SH3 repeats are not dependent on the inter-SH3 length, variations in inter-SH3 length may still alter the force-dependent energy dependency of the binding of dual-SH3 repeats to the vinculin linker. This occurs by adding a conformational constraint – under force, the binding of a shorter inter-SH3 length result in a less extended vinculin linker between the bound SH3 domains, while that of a longer inter-SH3 length can accommodate more extended vinculin linker conformations. A less extended vinculin linker in the bound complex leads to force-dependent decrease in the affinity of dual-SH3 domains with shorter inter-SH3 lengths. This is because upon dissociation, it leads to a longer extension change of the unbound vinculin linker in the direction of the applied force that would result in a reduction of free energy [14]. To investigate this further, we developed a model of this interaction.

### Modelling force-dependent binding of dual-SH3 repeats to the vinculin linker region

To investigate the potential force dependence of binding, we first examined the binding configurations of the 26 amino acid vinculin linker region (<u>ERAPPRPPLPREGLAPVRPPPPET</u>DD) containing the shared core binding sequence (underlined) with the dual-SH3 repeats. Next, we analyzed the force-dependent energies associated with each configuration. Finally, we derived a force-dependent binding constant based on these energies.

Figure 6 shows the four configurational states, unbound state denoted by “0”, two partially bound states with only one bound SH3 denoted by “1a” and “1b”, and a fully bound state with two SH3 domains bound denoted by “2” (Fig. 6a). Each of these configurational states is associated with a characteristic force-extension curve *x*_*i*_(*f*), and a force-dependent free energy term 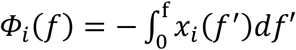, where *f* denotes the force and *i* denotes the state (Fig. 6b).

Analytical expressions of the force-extension curves for the four states were obtained with the following approximations: i) worm-like chain polymer model for the unbound polypeptide polymer with a bending persistence length of 0.8 nm for proline-rich sequences [48], suitable for state “0” and the unbound part in “1a” and “1b”; ii) the force-extension curve of a freely rotating rigid body of certain rigid body length, suitable for the bound region in states “1a” and “1b” with a length *d*_1_ ≈ 2.7 nm (Supp. Table 5 & Supp. Fig. 10), and that in state “2” with a rigid body length *d*_2_ (Fig. 6a). The values of *d*_1_ and *d*_2_ are estimated based on the AlphaFold2 predicted structures which were relaxed using full-atom molecular dynamics simulations. The values of *d*_2_ depend on the inter-SH3 length of the dual-SH3 model protein (Supp. Table 3 & Supp. Table 6).

Based on the four configurational state model, an analytical expression of the force-dependent dissociation constant can be derived:

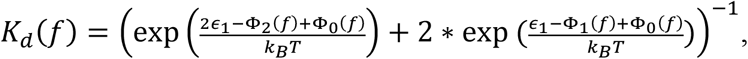

where −*ϵ*_1_ is the binding energy between a single SH3 and its binding site. Here we have assumed the two SH3 have similar binding energies for simplicity (Supp. Fig. 12 & Supp. Info. 3 for derivation).

Figure 6c shows the predicted *K*_*d*_(*f*)/*K*_*d*_(*f* = 0) curves for several *d*_2_ values from 4.2 - 5.0 nm and two single-SH3 binding energies of −10 *k*_*B*_*T* and −20 *k*_*B*_*T*, corresponding to a wide range of single-domain dissociation constant from 45 *μ*M to 2 nM. The results suggest a strong effect of the value of *d*_2_ on *K*_*d*_(*f*). At *d*_2_ values smaller than 5.0 nm, the model predicts that within ∼5 pN, increasing force has a weak enhancement of the binding affinity (a fold decrease of ∼ 1.6 in the Kd), whereas at forces above ∼5 pN, significant force-dependent suppression of the binding is observed. The dissociation constant at >10 pN is several folds greater than that at 0 pN. The curves obtained at the two single-SH3 binding energies overlap each other, indicating *ϵ*_1_ is an insensitive parameter of the model over the single-domain binding energy range (Fig. 6c, Supp. Info. 4 & Supp. Fig. 13).

Together, the results from theoretical modeling suggest a dual-layer mechanical regulation of the binding affinity between the vinculin linker and its binding partners. The physiological forces applied to full-length vinculin not only switch the vinculin linker from a less accessible closed conformation to a more accessible open conformation, but also regulate the binding affinity between the open conformation of the linker and its binding partners in a tension-dependent manner. The dual-SH3 model protein proposes a new mode of mechanosensing, where the force-dependent binding affinity is influenced by the inter-SH3 length. A strong suppressive effect on the binding of dual-SH3 repeats is predicted at forces above 10 pN.

## III. DISCUSSION

Current models of vinculin show it as a mechanically inert linker that couples talin/alpha catenin to the actin cytoskeleton. In this study, we quantified the mechanical response of vinculin, and demonstrate that vinculin is significantly more complex mechanically than previously thought. We measured the mechanical stability of the force-bearing domains of vinculin when they are exposed to increasing force at loading rates of 0.4 and 4.0 pN per second. The results indicate that the vinculin helical bundle domains show switch-like behavior and unfold over a range of forces from a few pN to approximately 15 pN at loading rates in the order of pN/s, depending on each domain’s mechanical stability. This level of loading rate is physiologically relevant as the magnitude of tension experienced by vinculin is in the order of a few pN as estimated from the rate of retrograde actin flow (in the order of 10 nm/s [49, 50]). The mechanical stability of the vinculin domains is similar to that of the force-bearing domains in talin or α-catenin at similar loading rates [15, 51].

Our data show that the structural domains of vinculin can withstand tension within a range of a few pN up to 15 pN, at physiological force loading rates. This means that vinculin can buffer tension within this range until all its force-bearing domains are unfolded, corresponding to an extension of up to 150 nm (Fig. 7 and Supp. Info 2). This buffered tension range aligns with the estimated pN tension range measured using FRET tension sensors in live cells [52]. Our previous study also indicated that the tension-dependent structural transitions of talin rod α-helical bundles can buffer the talin tension over a similar range, consistent with the talin tension range reported with FRET tension sensors in live cells [32]. This suggests that the mechanical unfolding of force-bearing domains can provide an explanation for the physiological tension range for both vinculin and talin in live cells.

In our previous study examining the interactions between full-length vinculin and mechanically exposed VBS, we found that vinculin’s autoinhibition is relatively dynamic. This observation is supported by the high binding rate (on the order of 10^6^ M^−1^ s^−1^) and strong binding affinity, as indicated by a dissociation constant (*K*_d_) on the mechanically exposed VBS of approximately 12 nM [14]. In agreement with these findings, our current study did not reveal a mechanically stable head-tail association, which would have resulted in an additional stepwise extension increase of at least 10 nm at forces of a few pN (Supp. Info. 5, Supp. Fig. 14-15).

### Mechanical regulation of interactions with vinculin

Our findings indicate that the binding of vinculin to its partners is regulated by complex mechanical processes. At the domain level, the switch-like behavior of the vinculin 4-helix bundles will displace any proteins that engage the folded conformation, and recruit ligands that bind to the open conformation. Whilst the full interactome of vinculin is not known, it is likely that there are ligands for these domains in the different states that are recruited as a function of the mechanical signals and the resultant vinculin conformations generated.

Previous research has shown that most vinculin factors identified so far bind to the unstructured linker region between the head and the tail. Previous theoretical studies have predicted that binding-induced changes of the conformation and stiffness of a flexible binding site can lead to a significant change of binding affinity [48-50]. Therefore, forces applied to the flexible vinculin linker could have a significant impact on the binding of its binding factors. Indeed, using a dual-SH3 vinculin binding factor derived from CAP as a model, our theoretical modeling suggests that binding to the vinculin linker is optimized at few pN forces and is strongly suppressed at forces greater than 10 pN. This suppression can be explained by the deformation of the binding site caused by the binding which shortens the binding site, which opposes the force-dependent extension of the binding site. Overall, these results, in conjunction with previous studies that have shown that vinculin linker binding factors are suppressed from binding to full-length vinculin compared to isolated vinculin linker [38, 53, 54], suggest that there is a two-layer mechanical regulation of the vinculin linker binding factors - the release of the auto-inhibitory conformation of vinculin via binding to mechanically exposed VBS in talin or α-catenin, and the subsequent force-dependent interactions between the vinculin linker and its binding factors.

### The changing dimensions of a vinculin linkage

Another observation from this analysis of the switch-like behavior of the vinculin force-bearing domains is that each unfolding event has a large effect on the dimensions of the vinculin molecule. The complete unfolding of the helical bundles in D2-D4 and the linker results in a large extension change (Fig. 2b) and a total of 526 a.a. of peptide polymer will be released under tension. Considering that all the structural domains unfold over tension range from a few pN up to 15 pN at physiologically relevant stretching conditions, the result implies that vinculin tension-bearing domains can buffer tension within this force range during an extension change up to 150 nm (Supp. Info. 2), when it mediates tension transmission from talin or *α*-catenin to cytoskeleton network. Our recent analysis visualizing talin molecules to scale [55], changing up to 400 nm at 15 pN, illustrated how these changes in length lead to dramatic reorganization of molecules engaged to these mechanical scaffolds and here we show that a similar extension is occurring on the vinculin linkages. Coupling of the large extension changes in talin and the connected vinculin molecules indicates a meshwork of talin and vinculin switches that is incredibly plastic and can adopt numerous different switch patterns and architectures as a result of the mechanical signaling of the cell.

### Conclusion

In summary we show that vinculin has a complex mechanical response and has the potential to act as a mechanotransducer as well as a mechanoeffector. As with talin, the vinculin domains unfold at physiological forces, and the changes in dimensions of the linkages between talin and actin mediated by vinculin, and the proteins recruited to that site will be dependent on the mechanically induced conformations of each vinculin molecule.

## IV. METHODS AND MATERIALS

### Materials and Magnetic tweezers

The Halo-tagged Drosophila vinculin constructs (D3-D4, D2-D4, D1-D4 and D1-Vt) were made as described previously [32] using a custom vector comprising an N-terminal GST-tag for protein purification, which was cleaved using TEV protease, to yield vinculin proteins with N-terminal AVI-tag and C-terminal Halo-tags. An in house-made magnetic tweezers apparatus was used in this study with the paramagnetic beads (Cat.# AP-25-10, Spherotech) attached with short dsDNA handles [32]. In the force scanning process, forces were incrementally increased with different scan rates.

### AlphaFold2 Protein Structure Prediction

The amino acid sequences of the full-length Drosophila vinculin (UniProtKB - O46037 VINC_DROME), its linker (a.a. 728-786) with the first two SH3 domain (a.a. 60-182) of dCAP-E (UniProtKB - Q966U7_DROME) and SH3 repeats with varied inter-SH3 lengths were input at https://colab.research.google.com/github/sokrypton/ColabFold/blob/main/beta/AlphaFold2_advanced.ipynb [33, 34] for the AlphaFold2 protein structure prediction, and the structural models were analyzed using PyMOL software (The PyMOL Molecular Graphics System, Version 2.0 Schrödinger, LLC.). The structural models were validated by comparison with the crystal structure of human full-length vinculin (PDB 1TR2)[10].

### Molecular Dynamics Simulation

Two forms of MD simulation were performed, one without a pulling force and the other with a constant pulling force on the vinculin linker. GROMACS 2019.6 [56, 57] was utilized with an initial protein complex of the interacting residues (Supp. Table 8) from Drosophila vinculin linker with SH3 repeats of 13 a.a. and 25 a.a. inter-SH3 lengths predicted by the Alphafold2 [33, 34]. These simulations were performed under the AMBER99SB-ILDN force field [58] with the extended simple point charge (SPC/E) water model [59]. The initial complex was immersed in a periodic cubic box filled with 0.15 M NaCl solution. The steepest descents minimization of 50000 steps followed by a 100 ps-NPT and a 100 ps-NVT (300 K) equilibrations were performed. Afterwards, a 100 ns MD simulation was performed with its system coordinates saved every 10 *p* for further analysis. In the case where a pulling force was involved in the MD simulation, a constant force of 20 pN was applied to the vinculin linker, with its C-terminal residue fixed and the other end under the corresponding constant force.

## V. AUTHOR CONTRIBUTIONS

J.Y., B.G and N.B. conceived the research. X.L., Y.W. and M.Y., performed the experiments. K.B and B.K generated the reagents and stretchable protein fragments. X.L., and Y.W. analyzed the data. X.L., B.G. and J.Y. wrote the manuscript with help from all authors. J.Y. and B.G. supervised the research.

## VI. ACKNOWLEDGEMENT

This work is funded by the National Research Foundation (NRF), Prime Minister’s Office, Singapore under its NRF Investigatorship Programme (NRF Investigatorship Award No. NRF-NRFI2016-03) (to J.Y.), and grants from the National Research Foundation through the Mechanobiology Institute Singapore (to J.Y), BBSRC (BB/S007245/1 to B.G. and BB/S007318/1 to N.H.B.) and Cancer Research UK (CRUK-A21671 to B.G.),

## REFERENCES

1. Michael, M. and M. Parsons, New perspectives on integrin-dependent adhesions. Curr Opin Cell Biol, 2020. 63: p. 31–37.

2. Angulo-Urarte, A., T. van der Wal, and S. Huveneers, Cell-cell junctions as sensors and transducers of mechanical forces. Biochim Biophys Acta Biomembr, 2020. 1862(9): p. 183316.

3. Goult, B.T., N.H. Brown, and M.A. Schwartz, Talin in mechanotransduction and mechanomemory at a glance. J Cell Sci, 2021. 134(20).

4. Goldmann, W.H., Role of vinculin in cellular mechanotransduction. Cell Biol Int, 2016. 40(3): p. 241–56.

5. Klapholz, B. and N.H. Brown, Talin - the master of integrin adhesions. J Cell Sci, 2017. 130(15): p. 2435–2446.

6. Bays, J.L. and K.A. DeMali, Vinculin in cell-cell and cell-matrix adhesions. Cell Mol Life Sci, 2017. 74(16): p. 2999–3009.

7. Atherton, P., et al., Mechanosensitive components of integrin adhesions: Role of vinculin. Exp Cell Res, 2016. 343(1): p. 21–27.

8. Janostiak, R., et al., CAS directly interacts with vinculin to control mechanosensing and focal adhesion dynamics. Cell Mol Life Sci, 2014. 71(4): p. 727–44.

9. Goult, B.T., et al., RIAM and vinculin binding to talin are mutually exclusive and regulate adhesion assembly and turnover. J Biol Chem, 2013. 288(12): p. 8238–8249.

10. Borgon, R.A., et al., Crystal structure of human vinculin. Structure, 2004. 12(7): p. 1189–97.

11. Bakolitsa, C., et al., Structural basis for vinculin activation at sites of cell adhesion. Nature, 2004. 430(6999): p. 583–6.

12. Johnson, R.P. and S.W. Craig, F-actin binding site masked by the intramolecular association of vinculin head and tail domains. Nature, 1995. 373(6511): p. 261–4.

13. Chorev, D.S., et al., Conformational states during vinculin unlocking differentially regulate focal adhesion properties. Sci Rep, 2018. 8(1): p. 2693.

14. Wang, Y., et al., Force-Dependent Interactions between Talin and Full-Length Vinculin. J Am Chem Soc, 2021. 143(36): p. 14726–14737.

15. Yao, M., et al., Mechanical activation of vinculin binding to talin locks talin in an unfolded conformation. Sci Rep, 2014. 4: p. 4610.

16. Pang, S.M., et al., Mechanical stability of alphaT-catenin and its activation by force for vinculin binding. Mol Biol Cell, 2019. 30(16): p. 1930–1937.

17. Dumbauld, D.W., et al., How vinculin regulates force transmission. Proc Natl Acad Sci U S A, 2013. 110(24): p. 9788–93.

18. Maartens, A.P., et al., Drosophila vinculin is more harmful when hyperactive than absent, and can circumvent integrin to form adhesion complexes. J Cell Sci, 2016. 129(23): p. 4354–4365.

19. Ezzell, R.M., et al., Vinculin promotes cell spreading by mechanically coupling integrins to the cytoskeleton. Exp Cell Res, 1997. 231(1): p. 14–26.

20. Alenghat, F.J., et al., Analysis of cell mechanics in single vinculin-deficient cells using a magnetic tweezer. Biochem Biophys Res Commun, 2000. 277(1): p. 93–9.

21. Humphries, J.D., et al., Vinculin controls focal adhesion formation by direct interactions with talin and actin. J Cell Biol, 2007. 179(5): p. 1043–57.

22. Le Clainche, C., et al., Vinculin is a dually regulated actin filament barbed end-capping and side-binding protein. J Biol Chem, 2010. 285(30): p. 23420–32.

23. Thievessen, I., et al., Vinculin-actin interaction couples actin retrograde flow to focal adhesions, but is dispensable for focal adhesion growth. J Cell Biol, 2013. 202(1): p. 163–77.

24. Case, L.B., et al., Molecular mechanism of vinculin activation and nanoscale spatial organization in focal adhesions. Nat Cell Biol, 2015. 17(7): p. 880–92.

25. Hazan, R.B., et al., Vinculin is associated with the E-cadherin adhesion complex. J Biol Chem, 1997. 272(51): p. 32448–53.

26. Weiss, E.E., et al., Vinculin is part of the cadherin-catenin junctional complex: complex formation between alpha-catenin and vinculin. J Cell Biol, 1998. 141(3): p. 755–64.

27. Goldmann, W.H., et al., Vinculin, cell mechanics and tumour cell invasion. Cell Biol Int, 2013. 37(5): p. 397–405.

28. Biswas, R., et al., Mechanical instability of adherens junctions overrides intrinsic quiescence of hair follicle stem cells. Dev Cell, 2021. 56(6): p. 761–780 e7.

29. Kanoldt, V., et al., Metavinculin modulates force transduction in cell adhesion sites. Nat Commun, 2020. 11(1): p. 6403.

30. Vasile, V.C., et al., Identification of a metavinculin missense mutation, R975W, associated with both hypertrophic and dilated cardiomyopathy. Mol Genet Metab, 2006. 87(2): p. 169–74.

31. Maeda, M., et al., Dilated cardiomyopathy associated with deficiency of the cytoskeletal protein metavinculin. Circulation, 1997. 95(1): p. 17–20.

32. Yao, M., et al., The mechanical response of talin. Nat Commun, 2016. 7: p. 11966.

33. Senior, A.W., et al., Improved protein structure prediction using potentials from deep learning. Nature, 2020. 577(7792): p. 706–710.

34. Jumper, J., et al., Highly accurate protein structure prediction with AlphaFold. Nature, 2021. 596(7873): p. 583–589.

35. Stec, D.L. and B. Stec, Complete Model of Vinculin Suggests the Mechanism of Activation by Helical Super-Bundle Unfurling. Protein J, 2022. 41(1): p. 55–70.

36. Miller, P.W., et al., Analysis of a vinculin homolog in a sponge (phylum Porifera) reveals that vertebrate-like cell adhesions emerged early in animal evolution. J Biol Chem, 2018. 293(30): p. 11674–11686.

37. Ziegler, W.H., R.C. Liddington, and D.R. Critchley, The structure and regulation of vinculin. Trends Cell Biol, 2006. 16(9): p. 453–60.

38. DeMali, K.A., C.A. Barlow, and K. Burridge, Recruitment of the Arp2/3 complex to vinculin: coupling membrane protrusion to matrix adhesion. J Cell Biol, 2002. 159(5): p. 881–91.

39. Kioka, N., et al., Vinexin: a novel vinculin-binding protein with multiple SH3 domains enhances actin cytoskeletal organization. J Cell Biol, 1999. 144(1): p. 59–69.

40. Brindle, N.P., et al., The focal-adhesion vasodilator-stimulated phosphoprotein (VASP) binds to the proline-rich domain in vinculin. Biochem J, 1996. 318 (Pt 3): p. 753–7.

41. Choi, H.J., et al., alphaE-catenin is an autoinhibited molecule that coactivates vinculin. Proc Natl Acad Sci U S A, 2012. 109(22): p. 8576–81.

42. Chen, H., et al., Improved high-force magnetic tweezers for stretching and refolding of proteins and short DNA. Biophys J, 2011. 100(2): p. 517–23.

43. Zhao, X., et al., Studying the mechanical responses of proteins using magnetic tweezers. Nanotechnology, 2017. 28(41): p. 414002.

44. Wang, Y., J. Yan, and B.T. Goult, Force-Dependent Binding Constants. Biochemistry, 2019. 58(47): p. 4696–4709.

45. Mandai, K., et al., Ponsin/SH3P12: an l-afadin- and vinculin-binding protein localized at cell-cell and cell-matrix adherens junctions. J Cell Biol, 1999. 144(5): p. 1001–17.

46. Kioka, N., K. Ueda, and T. Amachi, Vinexin, CAP/ponsin, ArgBP2: a novel adaptor protein family regulating cytoskeletal organization and signal transduction. Cell Struct Funct, 2002. 27(1): p. 1–7.

47. Zhao, D., et al., Structural investigation of the interaction between the tandem SH3 domains of c-Cbl-associated protein and vinculin. J Struct Biol, 2014. 187(2): p. 194–205.

48. Yu, M., et al., Effects of Mechanical Stimuli on Profilin- and Formin-Mediated Actin Polymerization. Nano Lett, 2018. 18(8): p. 5239–5247.

49. Alexandrova, A.Y., et al., Comparative dynamics of retrograde actin flow and focal adhesions: formation of nascent adhesions triggers transition from fast to slow flow. PLoS One, 2008. 3(9): p. e3234.

50. Swaminathan, V., et al., Actin retrograde flow actively aligns and orients ligand-engaged integrins in focal adhesions. Proc Natl Acad Sci U S A, 2017. 114(40): p. 10648–10653.

51. Yao, M., et al., Force-dependent conformational switch of alpha-catenin controls vinculin binding. Nat Commun, 2014. 5: p. 4525.

52. Grashoff, C., et al., Measuring mechanical tension across vinculin reveals regulation of focal adhesion dynamics. Nature, 2010. 466(7303): p. 263–6.

53. Chen, H., et al., Spatial distribution and functional significance of activated vinculin in living cells. J Cell Biol, 2005. 169(3): p. 459–70.

54. Huttelmaier, S., et al., The interaction of the cell-contact proteins VASP and vinculin is regulated by phosphatidylinositol-4,5-bisphosphate. Curr Biol, 1998. 8(9): p. 479–88.

55. Barnett, S.F.H. and B.T. Goult, The MeshCODE to scale-visualising synaptic binary information. Front Cell Neurosci, 2022. 16: p. 1014629.

56. Mark James Abraham, T.M., Roland Schulz, Szilárd Páll, Jeremy C. Smith, Berk Hess, Erik Lindahl, GROMACS: High performance molecular simulations through multilevel parallelism from laptops to supercomputers. SoftwareX, 2015. 1-2: p. 19–25.

57. Tan, C., et al., Implementation of residue-level coarse-grained models in GENESIS for large-scale molecular dynamics simulations. PLoS Comput Biol, 2022. 18(4): p. e1009578.

58. Lindorff-Larsen, K., et al., Improved side-chain torsion potentials for the Amber ff99SB protein force field. Proteins, 2010. 78(8): p. 1950–8.

59. H. J. C. Berendsen, J.R.G., and T. P. Straatsma, The missing term in effective pair potentials. J. Phys. Chem, 1987. 91(24): p. 6269–6271.

