## Supplementary Information for "The mechanical response of vinculin"

**Supp. Info. 1 Converting the force-dependent step sizes into the contour length of the protein peptide released during unfolding.**

As an example, let's examine the unfolding of vinculin D3-D4. The D4 domain of vinculin features a single four-helix bundle, while the D3 domain has two four-helix bundles that share a long α-helix. It is believed that the α-helix bundles behave similarly to disordered peptides when they are unfolded. In previous studies, the contour length of a disordered peptide was calculated to be approximately 0.38 nm per amino acid residue [1, 2].

Considering the helical domains in vinculin D3-D4 contain 333 amino acid residues, the unfolding D3-D4 construct would result in release of polypeptide of a contour length of approximately $L = 126$ nm (calculated by multiplying 333 by 0.38). The force-dependent extension change upon unfolding can be calculated as $\Delta x(f) = x_p(f) - x_d(f)$, which is the difference between the force-dependent extension of the released polypeptide, $x_p(f)$, and that of the folded D3-D4 domains, $x_d(f)$.

$x_p(f)$ can be calculated based on worm-like chain polymer model using Marko-Siggia formula [3]:

$$\frac{fA}{k_BT} = \frac{x}{L} + \frac{1}{4*\left(1-\frac{x}{L}\right)^2} - \frac{1}{4} \qquad (1)$$

where $A$ is the persistence length of polypeptide in the range of 0.5 to 0.8 nm which depends on the amino acid sequence, and $L = \text{number of amino acids} \times 0.38$ nm [1, 2] is the contour length.

$x_d(f)$ can be calculated based on the force-extension curve of a rigid body that can rotate around an anchoring point. It can be modelled as a straight rod of a length equal to the linear distance $d$ between the two force-attaching points on the rigid body. $x_d(f)$ has a simple analytical expression that is identical to the force-extension curve of a single segment in freely-joint chain polymer model:

$$\frac{x_d}{d} = coth\frac{fd}{k_BT} - \frac{k_BT}{fd} \qquad (2)$$

Using $A = 0.8$ nm, the predicted contour lengths of unfolded domains are consistent with the number of amino acids in those folded structures.

When examining the 2D scatter plot for D3-D4 (as shown in the right panel of Fig. 3a), it can be observed that group A signals unfold at approximately 8 pN, and a contour length change of roughly 30 nm. On the other hand, group B signals unfold at 10-13 pN, which is associated with a contour length change of around 45 nm. In total, the contour length change for all three domains (vinculin D3-D4) is approximately 30 + 2 × 45, which is approximately 120 nm. This value is close to the estimated change in bead height (~126 nm).

**Supp. Info. 2 The extension of the fully unfolded polypeptide at 15 pN.**

The force-bearing domains of Drosophila vinculin span from D1B to D4 and are arranged as bundles of α-helices. When these regions become unfolded, the resulting polypeptide behaves as a disordered polypeptide polymer. Since there are 569 amino acid residues in the helical bundles of vinculin D1B-D4, the complete unfolding of these domains would result in a peptide contour length of 569 × 0.38 nm, or roughly 216 nm. By applying the Marko-Siggia formula [3] (Eq. 1) and a polymer persistence length of 0.8 nm, the extension of the completely unfolded polypeptide would be approximately 0.7 × 216, or about 151 nm, when subjected to a force of 15 pN.

**Supp. Info. 3 Determination of the force-dependent dissociation constant for the Drosophila vinculin linker with dual-SH3 repeats using a four-state model.**

A 26 a.a. vinculin linker segment is modelled with dual-SH3 repeats, represented in schematic illustrations (Fig. 6). The tandem interaction has four states: "0" for no binding, "1a" or "1b" for single binding, and "2" for dual binding. The OFF state indicates no binding, while the ON state denotes one of the binding sites is bound. The linker is a flexible polymer in state "0," a combination of a flexible polymer and a rigid bound region in state "1a/b," and adopts a rigid bound conformation in state "2." The linear distances between the force-attaching points on the rigid bound regions are denoted by $d_1$ and $d_2$ for states "1a/b" and "2," respectively (Supp. Fig. 13).

There are four possible states for the interaction between the drosophila vinculin linker and dual-SH3 model protein. The first is the "off" state, denoted by "0", where no SH3 domains bind to the vinculin linker. The second to the fourth states are the "on" states, denoted by "1a", "1b" and "2", where at least one SH3 domain binds to the linker. At equilibrium, based on Boltzmann distribution, the ratio of the probability of bound state to the unbound states is:

$$\frac{p_{on}(f)}{p_{off}(f)} = \frac{e^{-g_2(f)/k_BT} + 2 * e^{-g_1(f)/k_BT}}{e^{-g_0(f)/k_BT}} \quad (3)$$

where $g_i$ represents the free energy of state "i". $k_B$ is the Boltzmann constant, $T$ is the temperature in Kelvin.

Setting the zero-force unbound state as the reference, the force-dependent energies of the states can be written as:

$$g_0(f) = \Phi_0(f) \quad (4)$$

$$g_1(f) = \Phi_1(f) - \epsilon_1 - k_BTln(c) \quad (5)$$

$$g_2(f) = \Phi_2(f) - 2\epsilon_1 - k_BTln(c) \quad (6)$$

Where $\Phi_i(f) = -\int_0^f x_i(f')df'$ with $x_i(f)$ being the force-extension curve of the state "i", $\epsilon_1$ denotes the binding energy of a single SH3 domain on the linker; and $c$ is the ligand concentration in Molar units.

Based on $\frac{p_{on}(f)}{p_{off}(f)} = \frac{c}{K_d(f)}$, we have:

$$K_d(f) = c\frac{p_{off}(f)}{p_{on}(f)} = c\frac{e^{-g_0(f)/k_BT}}{e^{-g_2(f)/k_BT} + 2 * e^{-g_1(f)/k_BT}}$$

$$K_d(f) = \left(exp\left(\frac{2\epsilon_1 - \Phi_2(f) + \Phi_0(f)}{k_BT}\right) + 2 * exp\left(\frac{\epsilon_1 - \Phi_1(f) + \Phi_0(f)}{k_BT}\right)\right)^{-1} \quad (7)$$

Based on Eq. (7), the ratio of $K_d(f)$ to that at zero force $K_{d0}$ is the zero-force dissociation constant is:

$$\frac{K_d(f)}{K_{do}} = \frac{exp\left(\frac{2\epsilon_1}{k_BT}\right) + 2 * exp\left(\frac{\epsilon_1}{k_BT}\right)}{exp\left(\frac{2\epsilon_1 - \Phi_2(f) + \Phi_0(f)}{k_BT}\right) + 2 * exp\left(\frac{\epsilon_1 - \Phi_1(f) + \Phi_0(f)}{k_BT}\right)} \quad (8)$$

The force-extension curve $x_i(f)$ of each state is described as follows (Supp. Fig. 13):

- $x_0(f)$ for the flexible linker is calculated by the worm-like chain model of polypeptide chain with a bending persistence length $A$ typically over a range from 0.5 nm to 0.8 nm, using the Marko-Siggia formula (Eq. 1) [3], where $L = 26 \times 0.38$ nm is the contour length of the 26 a.a. that provides the binding sites for the dual-SH3.
- $x_1(f) = x_{1p}(f) + x_{1b}(f)$ for the peptide bound with one SH3 domain is calculated by the sum of the part ($x_{1b}(f) = \mathrm{d}_1(\coth\frac{fd_1}{k_BT} - \frac{k_BT}{fd_1})$) contributed from rotation of a rigid rod of a length of $d_1 = 2.56$ nm predicted by AlphaFold2 and the other part ($x_{1p}(f)$) from the remaining flexible peptide of 13 amino acids using the worm-like chain polymer model (Eq. 2).
- $x_2(f) = d_2(\coth\frac{fd_2}{k_BT} - \frac{k_BT}{fd_2})$ for the fully bound state is calculated from rotation of a rigid rod of a length of $d_2$ predicted by AlphaFold2.

Figure 6c in the main text plots $\frac{K_d(f)}{K_{d0}}$, where $K_{d0}$ is zero-force $K_d$ by setting $f = 0$ pN, with binding energies of $\epsilon_1$ of $10k_BT$ and $20k_BT$, respectively.

**Supp. Info. 4 The predicted $\frac{K_d(f)}{K_{d0}}$ is insensitive to the single-SH3 binding energy $\epsilon_1$ in the model.**

For $\epsilon_1 \gg k_BT$ and $\gg \Phi_0(f) - \Phi_1(f)$, it is easy to show from Eq. (8) that: $\frac{K_d(f)}{K_{d0}} \approx \exp\left(\frac{\Phi_2(f)-\Phi_0(f)}{k_BT}\right)$, which is independent on the value of $\epsilon_1$. This is the case for the values $\epsilon_1$ greater than $10k_BT$ over a force range from 0 to 20 pN. As a result, the curves for $\frac{K_d(f)}{K_{d0}}$ overlap, as shown in Fig. 6c.

However, for smaller values of $\epsilon_1$, for instance, $5k_BT$ (see Supp. Fig. 14), the $\frac{K_d(f)}{K_{d0}}$ curve overlaps with that of $\epsilon_1 = 20k_BT$ only for force values less than 10 pN. For force values greater than 10 pN, the $\frac{K_d(f)}{K_{d0}}$ curve starts to deviate from that of $20k_BT$.

**Supp. Info. 5 Estimating the extension change (in nm) of Drosophila vinculin during head-tail dissociation**

Based on the head-tail dissociated structure, approximately the full-length vinculin can be considered as a mixture of three rigid bodies and a 59 a.a. linker (Supp. Fig. 16). The force-extension curve can be estimated by:

$$x(f) = d_1 coth(\frac{fd_1}{k_BT} - \frac{k_BT}{fd_1}) + d_2 coth(\frac{fd_2}{k_BT} - \frac{k_BT}{fd_2}) + d_3 coth(\frac{fd_3}{k_BT} - \frac{k_BT}{fd_3}) + x_{linker} \quad (9)$$

According to this estimation (plotted as Supp. Fig. 15), at forces above 2 pN, the extension of the full-length vinculin with all structural domains folded can reach above 10 nm. At forces above 5 pN, it is more than 20 nm.

Therefore, in the presence of a mechanically stable head-tail association, one should observe step with a step size more than 10 nm at a few pN forces, in addition to the unfolding steps from the structural domains, which was not observed in our experiments.

#### References

1. Chen, H., et al., *Dynamics of equilibrium folding and unfolding transitions of titin immunoglobulin domain under constant forces.* J Am Chem Soc, 2015. **137**(10): p. 3540-6.
2. Winardhi, R.S., et al., *Probing Small Molecule Binding to Unfolded Polyprotein Based on its Elasticity and Refolding.* Biophys J, 2016. **111**(11): p. 2349-2357.
3. Marko, J.F. and E.D. Siggia, *Stretching DNA.* Macromolecules, 1995. **28**(26): p. 8759-8770.

### Supporting Figures

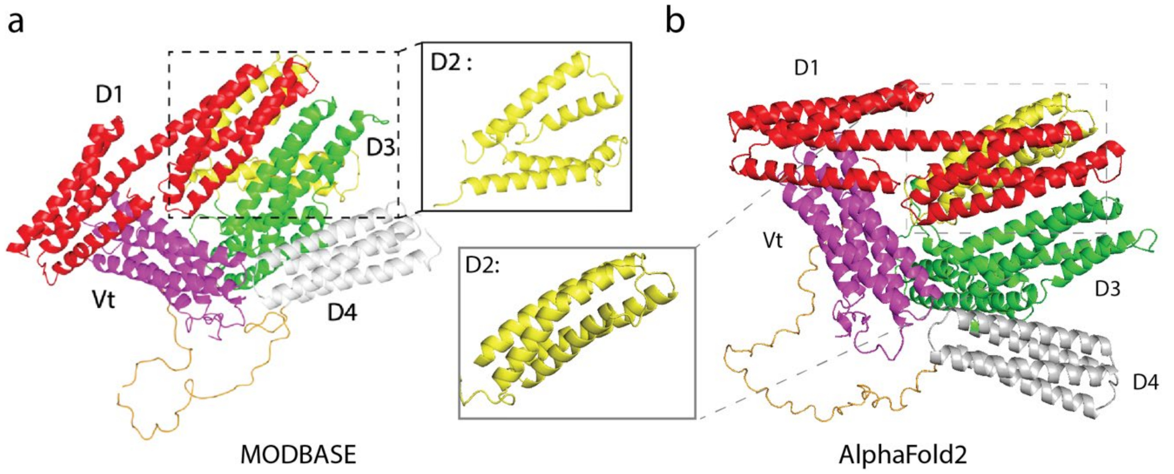


**Supp. Fig. 1 The protein structures of full-length Drosophila vinculin predicted by MODBASE (ref. no. O46037) (a) and AlphaFold2 (b).**

The prediction made by MODBASE was based on the chicken vinculin structure (PDB ID. 1ST6). The five domains of Drosophila vinculin (D1-D4 and Vt) are labelled, with an enlarged view of the D2 domain.

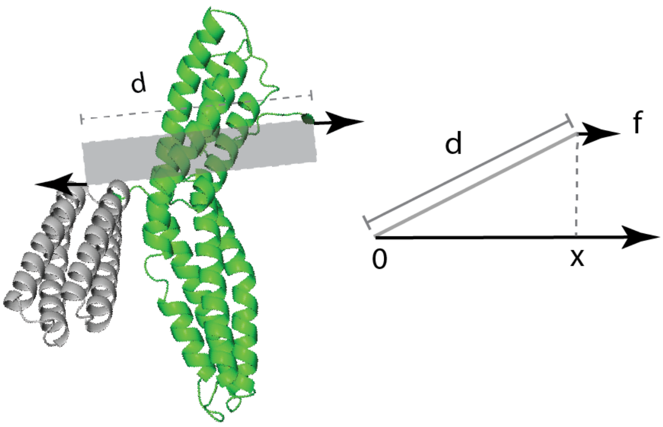


**Supp. Fig. 2 Illustration of Drosophila vinculin D3-D4 domains modelled as a rigid body with length d and under force $f$.**

The D3-D4 subdomains of Drosophila vinculin were modeled as a rigid body (shaded) in the unfolded state, with D3 shown in green and D4 in grey. Force $f$ was applied at the N-terminus of the D3 domain and the C-terminus of the D4 domain (indicated with arrows), resulting in an extension $x$ of the D3-D4 subdomains in the direction of the force (shown in the right panel).

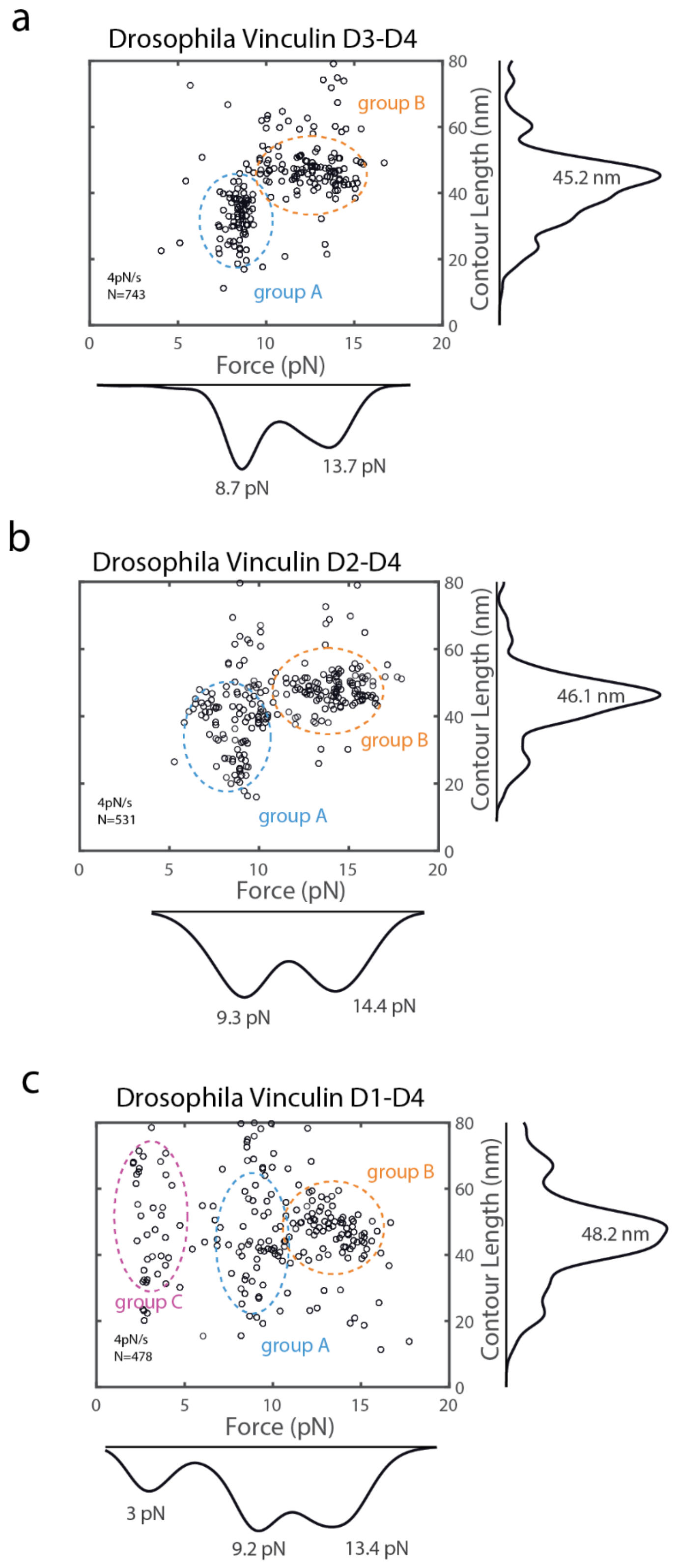


**Supp. Fig. 3 Mechanical responses of Drosophila vinculin at a different loading rate.**

The Drosophila vinculin sub-segments, including the D3-D4, D2-D4, and D1-D4 domains, underwent stretching at a loading rate of 4 pN/s. The resulting unfolding events were displayed on 2D scatter plots and grouped as A, B, and C based on the contour lengths of the released polypeptide polymer and the unfolding forces. Over 400 unfolding events from at least 3 independent tethers were used to obtain data points for each sub-segment.

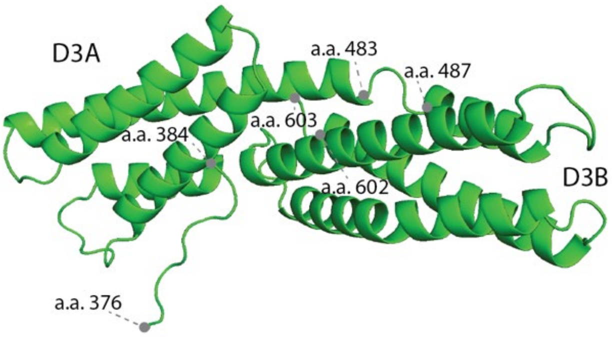


***Supp. Fig. 4 The D3 domain (amino acids 376-603) of Drosophila vinculin predicted using AlphaFold2.***

The AlphaFold2 prediction of Drosophila vinculin reveals helical bundle structures in the D3A subdomain spanning amino acids 384 to 483, and D3B subdomain spanning amino acids 487 to 602.

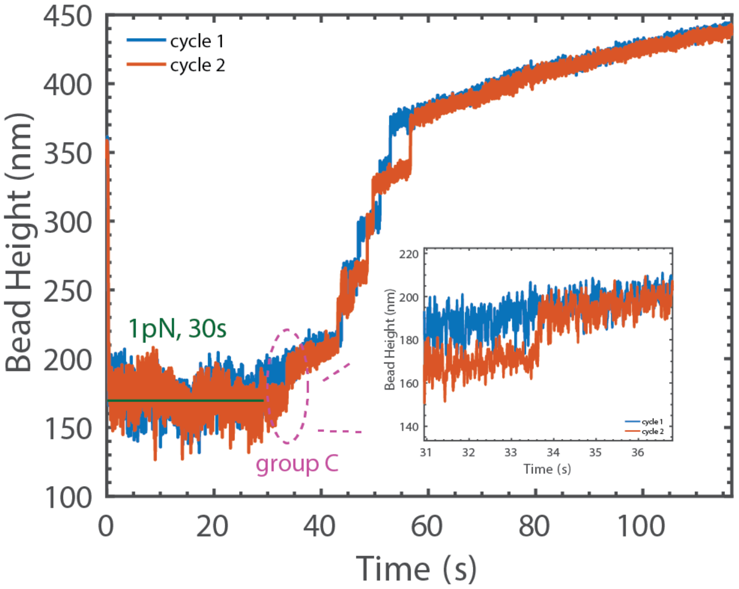


**Supp. Fig. 5 Bead height-Time plot of a vinculin D1-D4 subdomain.**

The smoothed data, obtained using the Savitzky-Golay smooth method in Matlab, for the vinculin sub-segment D1-D4 of Drosophila were plotted. The plot shows consecutive cycles of the sub-segment being held at 1 pN for 30 s, followed by being stretched at a loading rate of 0.4 pN/s. (Inset) The unfolding in group C was zoomed in to show that a step-wise unfolding was clear for cycle 2 (red) but not for cycle 1 (blue).

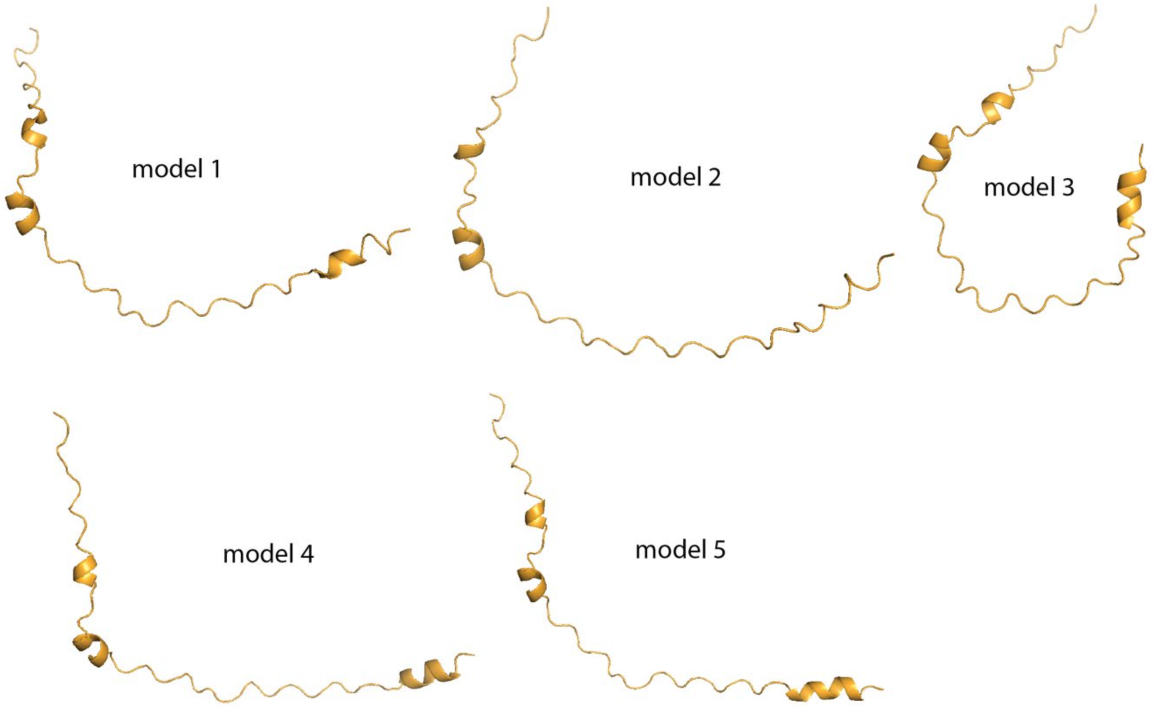


**Supp. Fig. 6 Predicted conformations of the isolated Drosophila vinculin linker generated using AlphaFold2.**

Five models of the isolated Drosophila vinculin linker predicted using AlphaFold2.

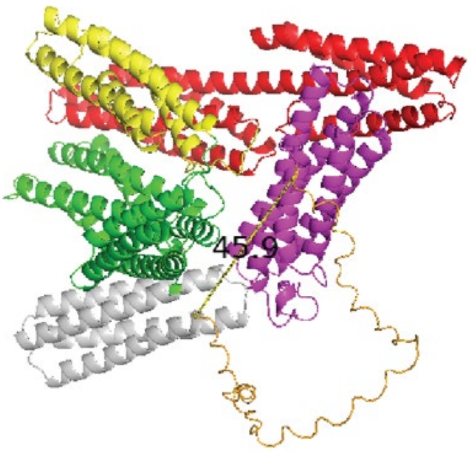


**Supp. Fig. 7 Measurement of the end-to-end distance of the linker.**

Full-length Drosophila vinculin (D1 in red, D2 in yellow, D3 in green, D4 in grey and Vt in purple) was predicted by AlphaFold2 and an end-to-end distance of its linker (gold) was measured to be 4.59 nm using PyMOL.

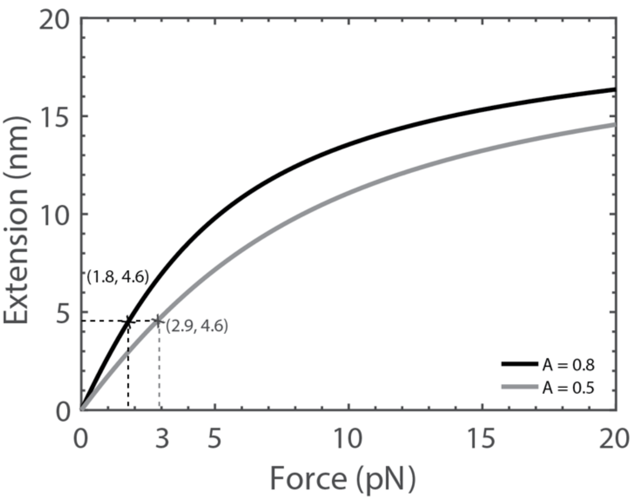


***Supp. Fig. 8 Force-extension curve of the vinculin linker.***

The linker consists of 59 amino acids and behaves similarly to an unstructured polypeptide polymer. Using the worm-like chain model, we obtained force-extension curves by applying polypeptide polymer persistence lengths (A) of 0.5 nm (in grey) and 0.8 nm (in black). For an extension of 4.6 nm in the linker (Supp. Table 2), either a force of 1.8 pN or 2.9 pN is resulted using A = 0.8 nm or 0.5 nm respectively.

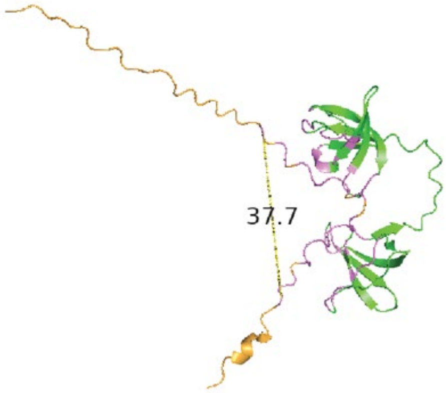


**Supp. Fig. 9 Measurement of the end-to-end distance of the interacting residues on the isolated vinculin linker.**

The isolated vinculin linker (in gold) interacts with a dual-SH3 repeat (in green) with an inter-SH3 length of 13 a.a. The predicted interacting residues are highlighted in purple. The end-to-end distance of 3.77 nm is measured between the first and the last residue of the interacting residues on the linker.

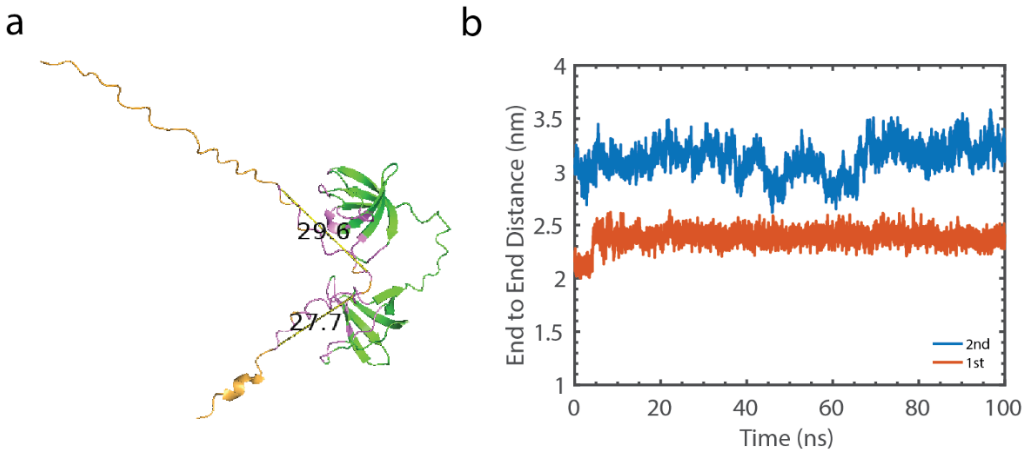


**Supp. Fig. 10 Measurement of the end-to-end distance of 1st and 2nd SH3-binding site on the vinculin linker.**

(a) The isolated vinculin linker (in gold) interacts with a dual-SH3 repeat (in green) with an inter-SH3 length of 13 a.a. The predicted interacting residues are highlighted in purple. The end-to-end distance of 2.77 nm is measured between residue 766 Proline (P) and residue 774 Threonine (T) of the 1st SH3-binding site on the linker, and that of 2.96 nm is measured between residue 751 glutamic acid (E) and residue 762 glutamic acid (E) for the 2nd SH3-binding site. (b) The end-to-end distance plot of 1st and 2nd SH3-binding sites upon MD simulation of 100 ns in the absence of external force. The 1st binding site maintains an average end-to-end distance of 2.4 nm, and that of 3 nm for the 2nd binding site.

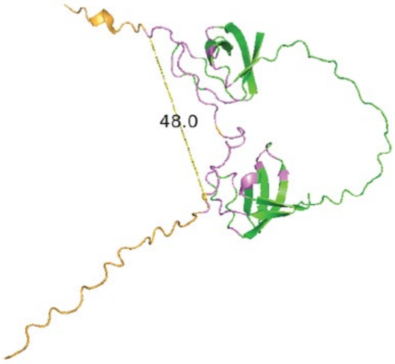


**Supp. Fig. 11 Measurement of the end-to-end distance of the interacting residues on the isolated vinculin linker.**

The isolated vinculin linker (in gold) interacts with a dual-SH3 repeat (in green) with an inter-SH3 length of 25 a.a. The predicted interacting residues are highlighted in purple. The end-to-end distance of 4.80 nm is measured between the first and the last residue of the interacting residues on the linker.

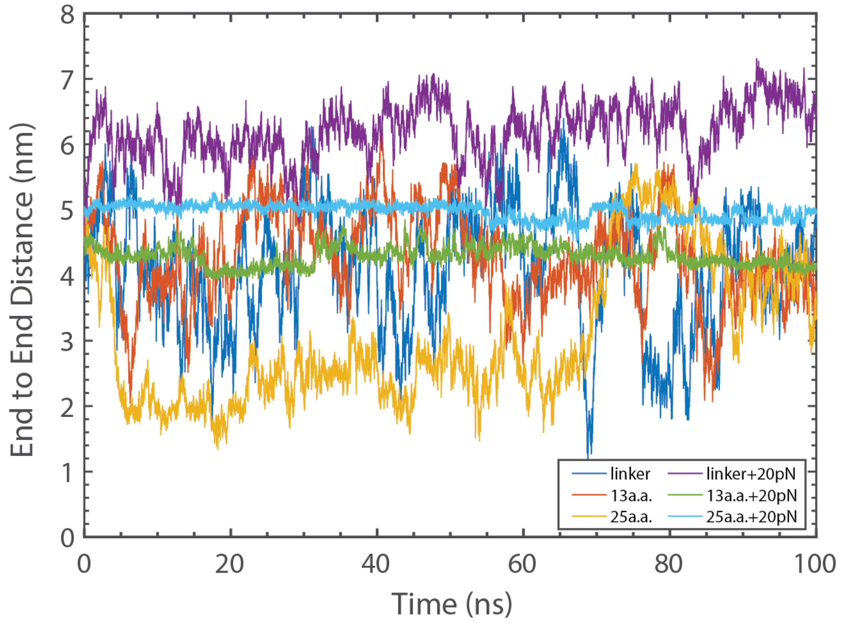


**Supp. Fig. 12 Plot of end-to-end distance of the binding site on the isolated Drosophila linker from MD simulations.**

The protein structures of the isolated vinculin linker or its complex with SH3 repeats, with various inter-SH3 linker lengths, were predicted and then subjected to a 100 ns MD simulation. The simulations were conducted with or without a constant force of 20 pN, and the end-to-end distances between the binding sites (which span from glutamic acid (E) at amino acid position 751 to Threonine (T) at amino acid position 774 on the vinculin linker) are presented.

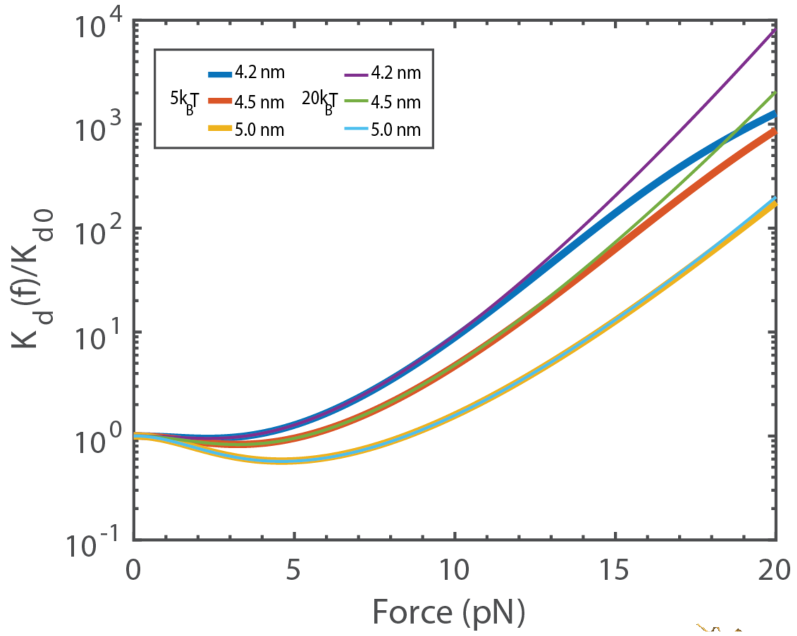


**Supp. Fig. 13 The $\frac{K_d(f)}{K_{d0}}$ plots.**

The graph plots the force-dependent dissociation constant $K_d(f)$ against $K_{d0}$, with a binding energy of $5k_BT$ and $20k_BT$ and $b_2$ values ranging from 4.2 to 5.0 nm, when $f = 0$.

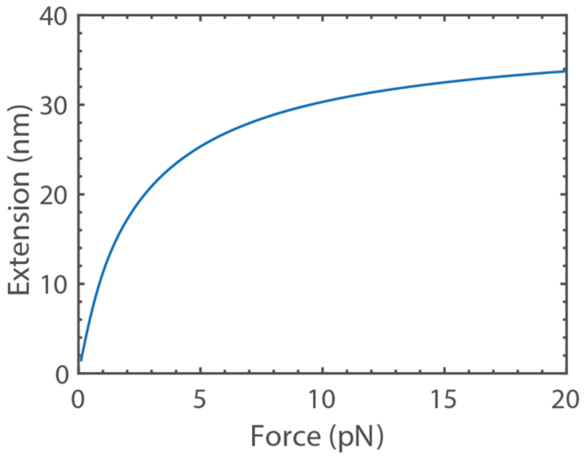


**Supp. Fig. 14 Force-extension curve of the full-length vinculin without domain unfolding.**

Eq. 9 was plotted with a polypeptide polymer persistence length (A) of 0.8 nm.

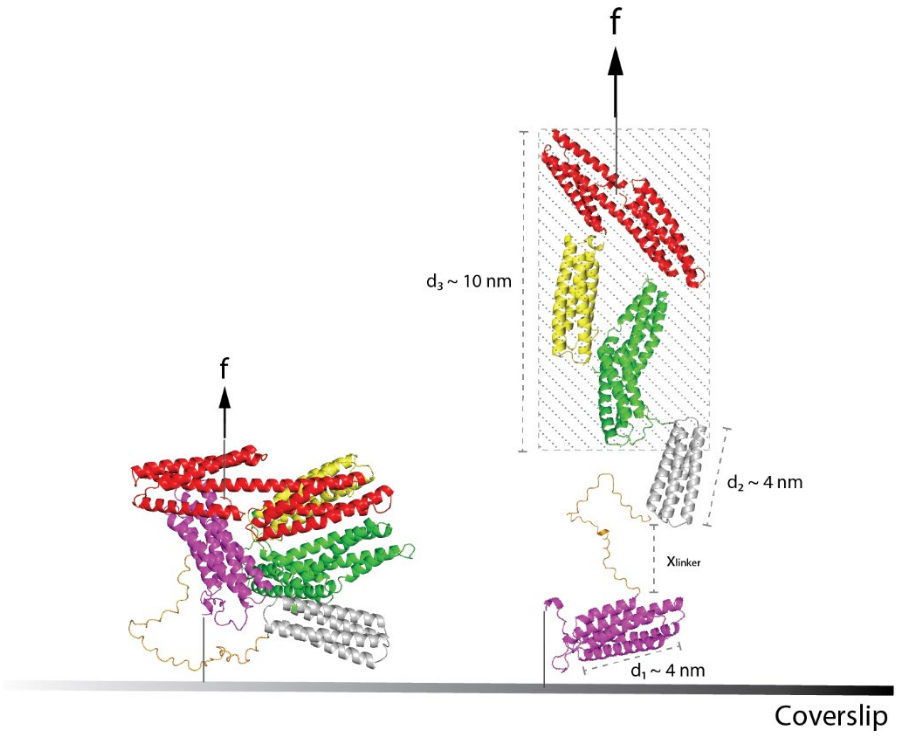


**Supp. Fig. 15 Illustration depicting the release of the head-tail association in Drosophila vinculin under the application of force (f).**

The height increase resulting from the helical bundles released from the head-tail association in Drosophila vinculin was measured using PyMOL. The analysis revealed a total increase of at least 10 nm in height at forces above 1 pN.

### Supporting Tables

**Supp. Table 1 The contour lengths and groupings of Drosophila vinculin domains**

| Drosophila Vinculin Domains | Helical Size (a.a.-a.a.) | Contour Length (CL) (nm) | Unfolding Forces | Grouping by CL and unfolding forces |
|---|---|---|---|---|
| D1 (D1A, D1B) | 241 (7 - 247) | ~ 91.58 | - | - |
| D2 | 116 (260 - 375) | ~ 44.08 | < 10 pN | group A |
| D3 subdomain A | 100 (384 - 483) | ~ 38 | < 10 pN | group A |
| D3 subdomain B | 116 (487 - 602) | ~ 44.08 | > 10 pN | group B |
| D4 | 117 (604 - 720) | ~ 44.46 | > 10 pN | group B |
| Vt | 150 (790 - 939) | ~ 57 | > 10 pN | - |

**Supp. Table 2 The end-to-end distance of the linker in the auto-inhibited full-length Drosophila vinculin measured using PyMOL.**

| Predicated outcomes by AlphaFold2 | Measured distance of vinculin linker in PyMOL (nm) |
|---|---|
| 1 | 4.55 |
| 2 | 4.59 |
| 3 | 4.63 |
| 4 | 4.65 |
| 5 | 4.59 |

**Supp. Table 3 The AlphaFold2-predicted interactions of the isolated Drosophila vinculin linker and the first two SH3 domains from dCAP-E**

| Predicted outcomes | Corresponding interacting residues on the linker | AlphaFold2 predicted distance between 1st and last a.a. of linker (nm) |
|---|---|---|
| 1 | (a.a. 751-776) ErAPPRPPLPREGlaPVRPPPPETDD | 5.48 |
| 2 | (a.a. 751-777) ERAPPRPPLPREgLApVrpPpPEtDdE | 1.69 |
| 3 | (a.a. 750-772) AEaAPPRPPLPREglaPVRPPPP | 5.35 |
| 4 | (a.a. 751-774) ERAPPRPPLPREglAPVRPPPPeT | 4.52 |
| 5 | (a.a. 751-776) ErAPpRPPLPrEgLAPVRPPPPeTDD | 3.77 |

* Interacting residues that are predicted are in capital letters.

**Supp. Table 4 The AlphaFold2-predicted residues on the isolated Drosophila vinculin linker that are involved in the interactions with dual-SH3 constructs of different inter-SH3 linker length (13 a.a., 19 a.a., 25 a.a. and 31 a.a.)**

| Predicted outcomes | 13 a.a. | 19 a.a. | 25 a.a. | 31 a.a. |
|---|---|---|---|---|
| 1 | ErAPPRPPLPREGlaPVRPPPPETDD | ErAPPRPPLPREGlAPVRPPPPETDD | ERAPPRPPLPREGlAPVRPPPPeT | ErAPPRPPLPREGlAPVRPPPPETDD |
| 2 | ERAPPRPPLPREgLApVrpPpPEtDdE | NaERAPPRPPLPrEgLAPVRPPPPETD | ErAPPRPPLPREGlAPVRPPPPETDD | ErAPPRPPLPREGLAPVRPPPPETDD |
| 3 | AErAPPRPPLPREglaPVRPPPP | NAERAPPRPPLPREglaPVRPPPPE | ERAPPRPPLPREglAPvRPPPPeT | ErAPpRPPLPrEGLAPVRPPPPeTDD |
| 4 | ERAPPRPPLPREglAPVRPPPPeT | ERAPPRPPLPRegLAPVRP | ErAPpRPPLPREgLAPVRPPPPeT | ERAPpRPPLPReglAPVRPPPPeT |
| 5 | ErAPpRPPLPrEgLAPVRPPPPeTDD | ErAPpRPPLPREGLAPVRPPPPeTDD | ErAPpRPPLPrEGLAPVRPPPPeTDD | ErAPpRPPLPREGlAPVRPPPPeTDD |

* Interacting residues that are predicted are in capital letters.

**Supp. Table 5 The AlphaFold2 predicted single-binding sites of the isolated Drosophila vinculin linker that are involved in the interactions with SH3 repeat of an inter-SH3 linker length of 13 a.a.**

| Single interacting SH3 domain | Corresponding interacting residues on the linker | AlphaFold2 predicted distance between 1st and last a.a. of linker (nm) |
|---|---|---|
| 2nd SH3 domain | (a.a 751-763) ErAPPRPPLPREG | 2.80 |
| | (a.a 750-762) AErAPPRPPLPRE | 2.94 |
| | (a.a 751-762) ERAPPRPPLPRE | 2.97 |
| | (a.a 751-762) ErAPpRPPLPrE | 2.96 |
| 1st SH3 domain | (a.a 766-776) PVRPPPPETDD | 2.19 |
| | (a.a 766-772) PVRPPPP | 1.69 |
| | (a.a 765-774) APVRPPPPeT | 2.12 |
| | (a.a 764-776) LAPVRPPPPeTDD | 2.77 |

* Interacting residues that are predicted are in capital letters.

**Supp. Table 6 The AlphaFold2 predicted dual-binding sites of the isolated Drosophila vinculin linker that are involved in the interactions with SH3 repeat of an inter-SH3 linker length of 25 a.a.**

| Inter-SH3 linker length | Corresponding interacting residues on the linker | AlphaFold2 predicted distance between 1st and last a.a. of linker (nm) |
|---|---|---|
| 25 a.a. | (a.a 751-776) ErAPPRPPLPREGlAPV RPPPPETDD | 4.80 |
| | (a.a 751-776) ErAPpRPPLPrEGLAPVR PPPPeTDD | 5.57 |
| | (a.a 751-774) ERAPPRPPLPREglAPvR PPPPeT | 5.42 |
| | (a.a 751-774) ErAPpRPPLPREgLAPV RPPPPeT | 4.02 |
| | (a.a 751-774) ERAPPRPPLPREGlAPV RPPPPeT | 5.15 |

* Interacting residues that are predicted are in capital letters.

### Supporting Movie

**Supp. Movie 1 Full-atom molecular dynamics simulation on the vinculin linker (highlighted in red) with both ends fixed (highlighted in cyan).**

The movie was captured at a frame rate of 0.3 frames per second for a 100 ns MD simulation.
